# Investigating the evolutionary plasticity of influenza virus splicing using influenza A virus segment 7 as a model system

**DOI:** 10.64898/2026.10.22.760983

**Authors:** Shu Zhou, Colin P. Sharp, Eleanor R. Gaunt, Rute Maria Pinto, Paul Digard

## Abstract

Influenza viruses A, C and D (IAV, ICV, IDV respectively) utilise mRNA splicing to produce two proteins from their matrix gene segment; the matrix (M1) protein and an ion channel (M2). In IAV, the M1 and M2 open reading frames (ORFs) partially overlap and splice donor (SD) and splice acceptor (SA) sites in the M1 gene are used to produce an M2 polypeptide with an N-terminal sequence shared with M1. In ICV and IDV, the M1 and M2 ORFs are concatenated and splice sites downstream of the M1 coding region are used to produce a matrix protein from the spliced mRNA. To probe how these different splicing strategies might have evolved, we used reverse genetics to test the ‘evolutionary’ flexibility of IAV SD and SA sites. We found that relatively small numbers of mutations were sufficient to activate a variety of alternative SD and SA sites in a mini-replicon system, several of which led to the production of ion channel isoforms with variant ectodomains, as well as one that produced an M1 variant. Furthermore, several of the viruses with altered splicing patterns were viable, including five with both SD and SA sites changed. Three of these ‘rewired’ viruses had only modest fitness penalties, despite up to 3 nonsynonymous changes in M1 and a 34 amino acid insertion in M2. These results suggest that IAV segment 7 splicing strategies are evolutionarily flexible and have implications for how IAV, ICV and IDV matrix segments might have diversified from a common ancestor.

## Introduction

Influenza viruses are enveloped viruses with segmented, negative-sense single-stranded RNA genomes that are classified into four species: A-D. Influenza A and B (IAV and IBV) have eight segments whereas influenza C and D (ICV and IDV) contain seven. At least nine essential viral proteins are encoded by these eight or seven segments. In all influenza viruses, the heterotrimeric PB1-PB2-PA/P3 (3P) RNA polymerase and nucleoprotein (NP) are encoded by segment 1-3 and 5, respectively. One copy of 3P and multiple copies of NP form a ribonucleoprotein particle (vRNP), which are the templates for transcription and replication of the viral genome (1-4). The glycoproteins vary between the influenza species: IAV and IBV segment 4 and 6 encode two spike proteins, haemagglutinin (HA) and neuraminidase (NA) (5), whereas ICV and IDV express a single hemagglutinin-esterase-fusion (HEF) glycoprotein, which combines HA and NA functions (6, 7). The matrix and nonstructural segments of all influenza viruses encode two essential viral proteins. The matrix protein (M1, BM1, CM1, DM1) and matrix 2 ion channel protein (M2, BM2, CM2, DM2) are translated from segment 7 in IAV and IBV and segment 6 in ICV and IDV, respectively (8-11). For all four species of influenza, M1 is the most abundant protein in the virion and serves as a structural scaffold (12, 13), while M2 works as an ion channel during viral entry (14-19). M2 is a relatively minor virion component; approximately 1000-fold lower than M1 in individual IAV viral particles (20). Nonstructural protein (NS1) and nuclear export protein (NEP) are encoded by segment 8 in IAV and IBV or segment 7 in ICV and IDV by alternative mRNA splicing (21, 22).

Genome sequence alignments among the four influenza viruses suggest that IDV clusters most closely with ICV, while IAV shows a closer evolutionary relationship with IBV; however, all four viruses share a common ancestor (10, 23). At the level of individual segments, the combination of nucleotide sequence alignments, protein structural and functional analyses indicate that the RNP components of polymerase and NP, HA/HEF, M1 and M2 and NS1/NEP (24-28) are conserved across all four species, indicating a common evolutionary origin. For the matrix gene products, AM1 and BM1 share primary sequence homology, as do CM1 and DM1, while the tertiary structure of the N-terminal domains of AM1 and CM1 indicate a common ancestry between all species (13). M2 proteins of all four species have similar topologies; small single pass transmembrane proteins with larger cytoplasmic domains than ectodomains (16). Structural and modelling studies indicate that IAV, IBV and ICV form similar left-handed 4-helix bundle structures (29, 30), consistent with a common ancestor. However, matrix gene expression strategies differ markedly between influenza A, B and C and D viruses. IBV expresses BM1 and BM2 proteins by translational termination-reinitiation (31), while IAV, ICV and IDV utilize two distinct strategies of alternative mRNA splicing (Figure 1A and B). In IAV, the unspliced mRNA is translated to produce M1 protein, while splice donor (SD) and splice acceptor (SA) sites near the 5’- and 3’-ends of the M1 gene are used to generate the mRNA2 spliced transcript, which encodes M2 (32, 33). Consequently, M1 and M2 share a common 9 amino-acid sequence at their N-terminus. Conversely, in ICV and IDV, the M1 and M2 ORFs share a longer concatenated ORF (Figure 1A) and neither SA site is within the mature M1 coding region. Instead, splice sites just upstream and within the ion channel coding region are utilised to generate M1 mRNA, either by forming a stop codon at the exon-exon junction (ICV) or by splicing into a very short second exon (IDV). The unspliced mRNA encodes a polyprotein that is thought to be post-translationally cleaved by signal peptidase to produce CM2/ DM2 (8-11, 34). Thus in contrast to the NS segment, protein function seems to have been conserved across the known influenza viruses while expression mechanisms diversified.

**Figure 1.**
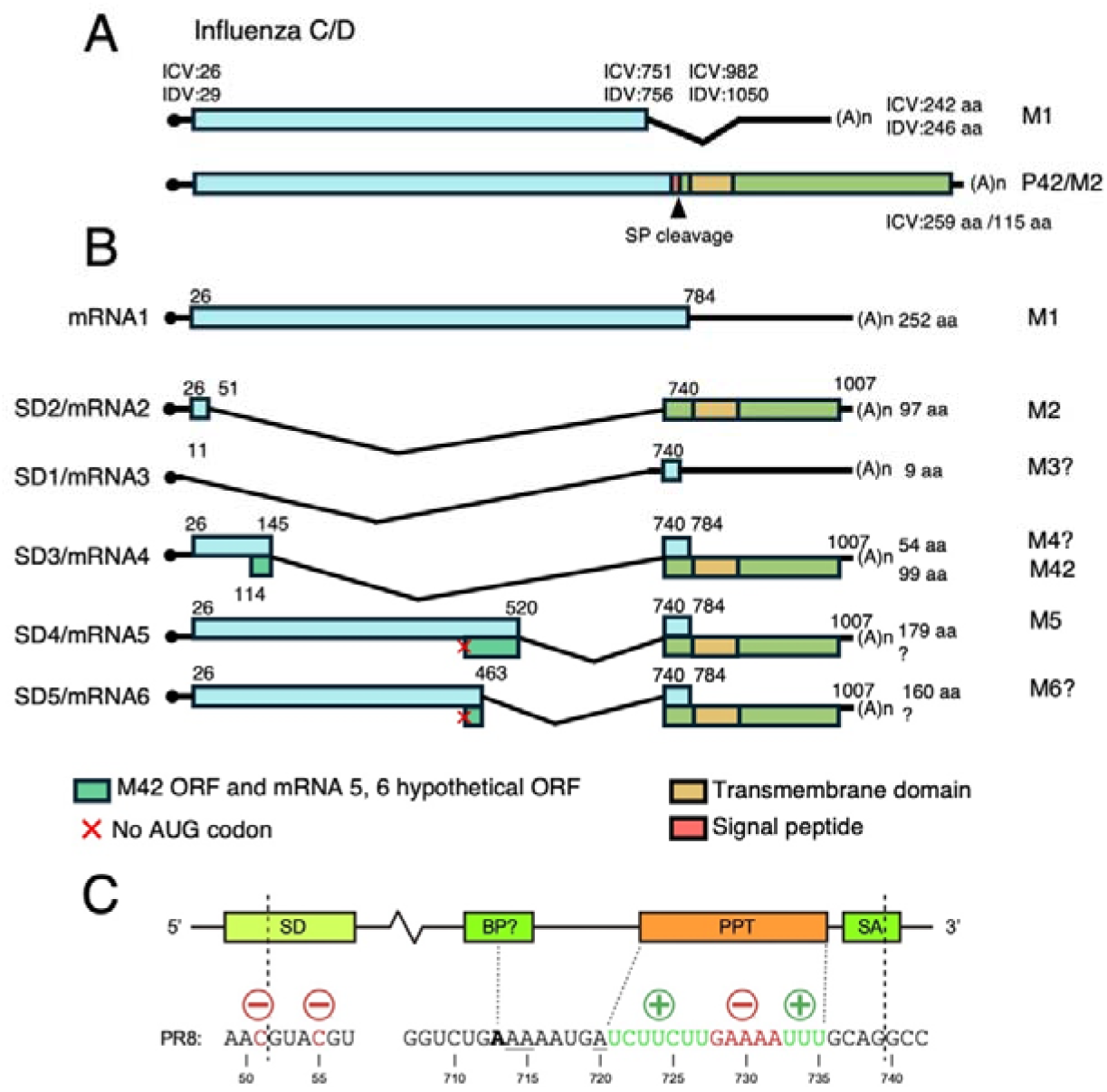
Influenza virus matrix gene transcript and ORF structures. Diagrammatic summary of (A) ICV/IDV segment 6 and (B) IAV segment 7 mRNA splicing strategies, drawn approximately to scale. Transcripts are listed in approximate order of abundance (mRNAs 1-3) and then numbers of virus strains known to produce them (mRNAs 4-6). The nucleotide locations of SD and SA sites are shown (10, 11, 47, 48). Potential ORFs are colour coded (blue: CM1/DM1/M1; green: CM2/DM2/M2; light greens: M42 ORF, mRNA5 and 6 hypothetical ORF) and total sizes (codons/ aa) are given on the right. Signal peptide and transmembrane domains (18, 47) are shown in red and yellow, respectively. (C) Splicing signals on IAV segment 7 mRNA. Segment 7 has a medium-strong splice donor (SD) and strong branch point (BP). The polypyrimidine tract (PPT) is interrupted by a purine-rich region, maintaining relatively weak splicing. A strong splice acceptor (SA) is located at the 3’ end of the intron (42).

Four conserved *cis*-acting sequence elements are required for pre-mRNA splicing: a SD motif at the 5’-end of the intron/ exon boundary; and three separate but usually juxtaposed elements: a branch point (BP), polypyrimidine tract (PPT) and 3’end of the intron and a 3’-SA sequence overlapping the intron/ exon boundary (Figure 1C). The SD is initially recognised by the U1 snRNP through complementary base pairing between the SD sequence and the RNA component of U1 snRNP (35, 36). Accordingly, SDs usually fall within a consensus sequence: MAG|GURUGU (35, 37). After cleavage at the 5’ end of the intron, the BP adenine (usually located in a loose consensus of YUN**<u>A</u>**Y) conjugates a guanine at the 5’-end of the intron to form a lariat structure (38). The heterodimer factor U2AF recognises PPT and SA by its two subunits (U2AF65 and U2AF35), respectively (36, 39). The 3’ SA is typically defined by the consensus sequence YAG, to facilitate recognition by U2AF35 (40), while the PPT is recognised by U2AF65 (41). The 5’-end of the cleaved SD and 3’-end of the SA are then ligated together, while the branch intron is degraded. Thus, in sum, an efficient splice unit requires the gain of four correctly-spaced functional motifs totalling around 30 ntds. IAV segment 7 splice signals generally, though imperfectly, resemble those of host cells to maintain the ratio of unspliced and spliced mRNAs needed to produce two essential viral proteins (32). Our previous study suggests that segment 7 has sub-optimal SD and PPT upstream of SA that set this balance (42) (Figure 1C).

Variation in the splicing pattern of evolutionary-related cellular genes is widespread eukaryotic lineages. Two theories about how alternative splicing patterns evolve from an ancestral gene have been proposed. In the sequence-based model, relaxed recognition of splice site motifs allows gradual mutational shifts between constitutive and alternative patterns (43, 44). Alternatively (or additionally), co-evolution of the splicing machinery and the RNA sequences it recognizes might have driven diversification of gene splicing at a deeper evolutionary level (43, 45). Regarding the closely related influenza viruses, the *cis*-factor-based model seems more plausible, as the family of viruses infect overlapping hosts and homeothermic tetrapod spliceosomes are conserved (46).

To test whether these principles of alternative splicing evolution also apply to influenza viruses, in this study, we investigated IAV splice site mobility. We found that multiple potential SD and SA sequences in segment 7 could be mutationally activated and that viable viruses using alternative splicing patterns to produce variant M1 and M2 proteins could be produced. We conclude that IAV shows a considerable level of splicing plasticity, supporting the idea that splice site migration could have occurred during orthomyxovirus evolution.

## Materials and Methods

### Bioinformatics analysis

To assess the strength of SD and SA motifs in WT and mutant segment 7s, MaxEntScan was used (49). The SD and SA sequences of A/Puerto Rico/8/1934 (PR8) segment 7 sequence (GeneBank accession: EF467824) were used as inputs. The MaxEntScan Python script (https://github.com/kepbod/maxentpy) was run on Google Colab (https://colab.research.google.com/). The SD input used 9-mers of the GT dinucleotide flanked by three upstream and four downstream bases, while the SA input used 23-mers of the AG dinucleotides preceded by 18 upstream and three downstream residues. The SD and SA strengths were calculated by score5 and score3 functions and visualised as a heatmap using the seaborn package (Python). A diverging colour scale centred at zero was used to represent decreases or increases in predicted SD or SA strength.

### Cells, viruses and plasmids

Human embryonic kidney cells (HEK 293T) cells (ATCC) and Madin-Darby canine kidney (MDCK) cells were cultured according to standard procedures (50). Cells were routinely tested for mycoplasma contamination. Influenza A/PR/8/34 (PR8) Wild type (WT) and modified viruses were rescued by transfections of eight plasmids corresponding to the eight viral genome segments under bidirectional polI and polII promoters (51) into HEK 293T cells using Lipofectamine 2000 following the manufacturer’s instructions. The virus was further propagated in MDCK cells as previous described (50). A plasmid containing bi-directional promoters (“pDUAL”) without a viral segment insert was used as a negative control (50). All constructs were confirmed by Sanger sequencing (GENEWIZ). For protein and mRNA analysis in single-cycle infections, cells were typically infected at an MOI of 5 (unless specified otherwise; see figure legends), incubated for one hour and then cells were washed to remove input viruses and medium was replaced. Virus work was carried out at the Roslin Institute under biological risk assessments BARA 1011 and GMRA 1811, approved by the local University of Edinburgh biosafety committee and the UK Health and Safety Executive.

### Mini-replicon assay

HEK 293T cells were seeded in 24-wells for protein analysis or 12-well plates for mRNA measurements. Cells cultured in 24-well plates were transfected with 50 ng of each desired combination of IAV reverse genetics plasmids, whereas cells cultured in 12-well plates were transfected with 100 ng. Transfections were performed using 1 ul or 2 ul Lipofectamine 2000, respectively, according to the manufacturer’s instructions (Thermo Fisher Scientific). The plasmids expressed combinations of the PB2, PB1, PA, NP, NS and M segments from either the bidirectional pDUAL vector containing RNA polymerase I and II promoters (51). The empty pDUAL vector plasmid was used as a negative control by replacing the segment 7 plasmid. The medium was removed 48 hours after transfection, and cells were lysed using Laemmli buffer (20% glycerol, 2% Sodium Dodecyl Sulfate (SDS), 100 mM Dithiothreitol (DTT), 24 mM Tris pH 6.8, 0.016% bromophenol blue) for western blotting, or lysed for total RNA extraction as described below.

### RT-PCR and splicing analysis

Total RNA was extracted from virus-infected or plasmid-transfected MDCK or 293T cells respectively using RNeasy Mini Kits (QIAGEN) according to the manufacturer’s instructions. After DNA treatment using a TURBO DNA-free Kit (Thermo Fisher Scientific), cDNA was generated from 800 ng of RNA using Lunascript RT Master Mix Kit (New England Biolabs, E3025) with oligo dT (IDT) as a primer. The cDNA was amplified using PrimeStar Max Premix (Takara Bio Inc.). M segment PCR was performed with the following primers: forward primer: GAGTCTTCTAACCGAGGTC for mRNA2 SD mutational analysis or TTCTAACCGAGGTCGAAAC for the rest of the analyses. A common reverse primer:

CCACAGCACTCTGCTGTTCCT was used. Segment 7 mRNA1, mRNA2 and mRNA4 were distinguished based on the sizes of their PCR products: 939 or 934 bp for mRNA1, 251 bp or 246 bp for mRNA2 and either 345 bp or 340 bp for mRNA4, depending on the forward primer used. NP mRNA PCR was performed with forward primer: ACGGAAAGTGGATGAGAGAACTCAT and reverse primer: CAACCGACCCTCTCAATATGAGTGC. GAPDH mRNA PCR was performed with forward primer: TGAAGGTCGGAGTCAACGGATTTGGT and reverse primer: CATGTGGGCCATGAGGTCCACCAC (52). PCR conditions were 98 °C for 5 min, 18 (for GAPDH) or 23 (for segment 5) or 28-30 cycles (for segment 7, depending on experiment) of 98 °C for 10s, 55 °C for 15s and 68 °C for 10s. PCR products were separated in 2%-2.5% agarose gels stained with SYBR™ Safe DNA Gel Stain (Thermo Fisher) and imaged using an Odyssey Fc imaging system (Li-Cor Inc.). PCR products of spliced transcripts were purified by Monarch DNA Gel Extraction Kit (New England Biolabs) and analysed by Sanger sequencing (GENEWIZ) and visulised using SnapGene.

### Western blotting analysis

SDS-PAGE followed by western blotting was performed using home-made 20% or 4-20% gradient gels (BioRad) and the Trans-Blot Transfer System (BioRad) according to the manufacturer’s instructions. Nitrocellulose membranes with 0.2 µm pores were blocked with phosphate-buffered saline (PBS)/0.1% Tween20/5% skimmed milk at room temperature for one hour, washed with PBS/0.1% Tween20 and incubated with primary antibody diluted in blocking media. Following overnight incubation at 4 °C, membranes were washed with PBS/0.1% Tween20 and incubated with fluorophore-labelled secondary antibody diluted in blocking media for one hour at room temperature. Membranes were imaged using an Odyssey Fc imaging system (Li-Cor Inc.). Image acquisition and analysis were performed using Image Studio Lite software. Purchased monoclonal antibodies were against α-tubulin (Thermo Fisher Scientific, Clone YL1/2), influenza M2 (Abcam, Clone 14C2), and influenza NP (Abcam, Clone C43). Purchased polyclonal antibodies were against influenza M1 (GeneTex, GTX125928) and (MRC PPU Reagents, University of Dundee, DA185) and M2 (Thermo Fisher Scientific, PA5-32233). In-house polyclonal rabbit antisera anti-NP (antiserum A2915), anti-M1 (antiserum A2917, raised against full-length M1) and anti-NS1 (antiserum V29) have been previously described (53). Purchased secondary antibodies were IRDye 800CW anti-Mouse (Li-Cor Inc, 925-32212), IRDye 680RD anti-Rat (Li-Cor Inc, 926-68076) and IRDye 800CW anti-Rabbit (Li-Cor Inc, 926-32213). Densitometric analysis of western blots was done using the gel plot function in ImageJ 2 (54).

### Virus titration and plaque size quantification

To titre viruses, confluent monolayers of MDCK cells were infected with serial-diluted viral samples in the presence of viral growth medium (serum free DMEM containing 0.14% BSA and 1 µg/ml TPCK trypsin) and incubated for one hour, followed by addition of 1.2% Avicel (Dupont) overlay 1:1 (v/v). After 48 hours, plates were formalin fixed and stained with 0.1% toluidine blue (Sigma). To quantify plaque size, fixed plates were scanned in an Epson Perfection V750 Pro Scanner at a resolution of 360 dpi. Images were converted to grey scale, contrast was adjusted and the area of each plaque was determined in ImageJ 2 (54). 5-20 plaques were selected and measured per experiment and normalised to the values from WT virus.

### Virion RNA extraction and sequencing

Viral genomic RNA was extracted from 140 µl of MDCK-grown virus stocks (containing 10^7^-10^8^ PFU/ ml) using a QIAamp Viral RNA Kit (QIAGEN) and the addition of 5.6 µg of carrier RNA according to the manufacturer’s instructions, and DNase-treated using the TURBO DNA-free Kit (Thermo Fisher Scientific). Reverse transcription was performed using Superscript IV (Thermo Fisher Scientific) and a Uni 12 primer (55) that targets a conserved nucleotide sequence present on all genome segment termini (AGCAAAAGCAGG). 1 µl of the resulting cDNA was used as a template for PCR using PrimeSTAR GXL Premix (Takara Bio Inc). Forward and reverse primers for PCR were AGCAAAAGCAGGTAGATATTG and AGTAGAAACAAGGTAGTTTTTTAC respectively. PCR conditions were 98°C for 5 min, 35 cycles of 98°C for 10s, 55°C for 15s and 68°C for 2 min. PCR products were purified by Monarch DNA Gel Extraction Kit (New England Biolabs) or Monarch Spin PCR & DNA Cleanup Kit (New England Biolabs) and analysed by Sanger sequencing (GENEWIZ).

### Statistical Analyses

All experiments that were statistically analysed (using GraphPad Prism 11) were performed in at least biological triplicate. Statistical tests used an ordinary One-way ANOVA test followed by Dunnett’s multiple comparisons test against PR8 WT. ****= P <0.0001; ***= P <0.001; **= P <0.01; *= P <0.05; absence of significance symbols = P>0.05. Error bars indicate the standard deviation (protein expression analysis) or standard error of the mean (plaque size comparisons).

## Results

### Intrinsic strength of SD influences SD usage in segment 7 with multiple SDs

In cellular mRNAs, alternative splicing patterns may evolve due to the coexistence of multiple weak SDs and relaxed recognition by the spliceosome (43, 44). In IAV segment 7, several other SDs apart from the primary SD2 (SD1, SD3 and 4; Fig 1B) have been identified (32, 56-58), but their usage varies by virus strain and is often weak at best (48, 56, 58). These observations are consistent with the *cis* model of how alternative splice sites evolve and further suggests that SD site selection in segment 7 may also be evolutionarily plastic. Thus, we systematically altered the strength of these minor viral SD site sequences to test how SD migration might occur during orthomyxovirus evolution.

Splice site migration implies not just the gain of a new SD (or SA) motif but also the loss of the original one. This could occur via mutational inactivation of a redundant SD motif, or, as competition between multiple SDs has been confirmed for cellular mRNA splice sites (59), by selection for a stronger SD sequence that then overrides use of the original SD. Competition between SD2 and SD3 in IAV segment 7 has already been demonstrated (48, 59). Furthermore, because both mRNA2 and mRNA4 encode functional ion channels (M2 and M42, respectively; Figure 1B) (48), SD migration between these two sites would maintain virus viability and might better reflect natural evolution. Accordingly, a panel of mutants was designed to gradually increase the similarity of the mRNA4 SD to the cellular SD consensus (Figure 2A). Accordingly, MaxEnt estimations of splice site strength ranged from the WT value of -1.72 to 10.86 (Supplementary Figure 1A). To avoid confounding effects from non-synonymous mutations in the M1 gene or overlapping segment-specific RNA packaging signals (60), we utilised a transfection-based mini-replicon system to then assay mRNA splicing from WT and mutant segments (42). HEK 293T cells were co-transfected or mock-transfected by PR8 segments 1-3 and 5 (encoding PB2, PB1, PA and NP, respectively; 3PNP), 8 (encoding NS1) and the various segment 7s. The viral polymerase catalysed segment 7 vRNA transcription while NS1 contributed to segment 7 mRNA splicing, nuclear export and overall cellular translation (61-66).We then assayed segment 7 RNA synthesis and protein expression by RT-PCR and western blotting, respectively. The WT segment 7 generated a mixture of unspliced mRNA1 and spliced mRNA2, but no detectable mRNA4 (Figure 2E lane 2). As successive mutations were introduced into SD3 to increase its strength, mRNA4 synthesis became visible and increased in abundance (lanes 3-5). In parallel, mRNA2 production reduced until it became undetectable. M1 mRNA1 levels were highest when only mRNA2 was produced (lane 2) and were clearly lowest (*i.e.* overspliced) when SD3 became dominant (lane 5). To assess ion channel protein production, we used two anti-M2 antibodies with differing specificities to differentiate accumulation of the M2 and M42 proteins: the 14C2 antibody recognizes residues 4 to 17 of the PR8 M2 protein but does not bind M42 (48), while a polyclonal antibody (PA5-32233) that was raised against amino acids 11-97 of M2 recognises both M2 and M42 but is unable to distinguish them. The WT segment expressed appreciable quantities of both M1 and M2 polypeptides (Figure 2E). However, M2 expression levels reduced when SD3 strength was increased, while levels of overall ion channel production (detected by the polyclonal C-terminal antibody) remained more constant. Double staining demonstrated that only M42 was produced and M1 levels were markedly reduced in the strongest mRNA4 SD (SD3) mutant (merge channel; lane 6). Thus, viral ion channel expression levels were consistent with transcript abundance. Furthermore, a strong SD3 can outcompete a WT SD2 without needing any mutational inactivation of the latter.

**Figure 2.**
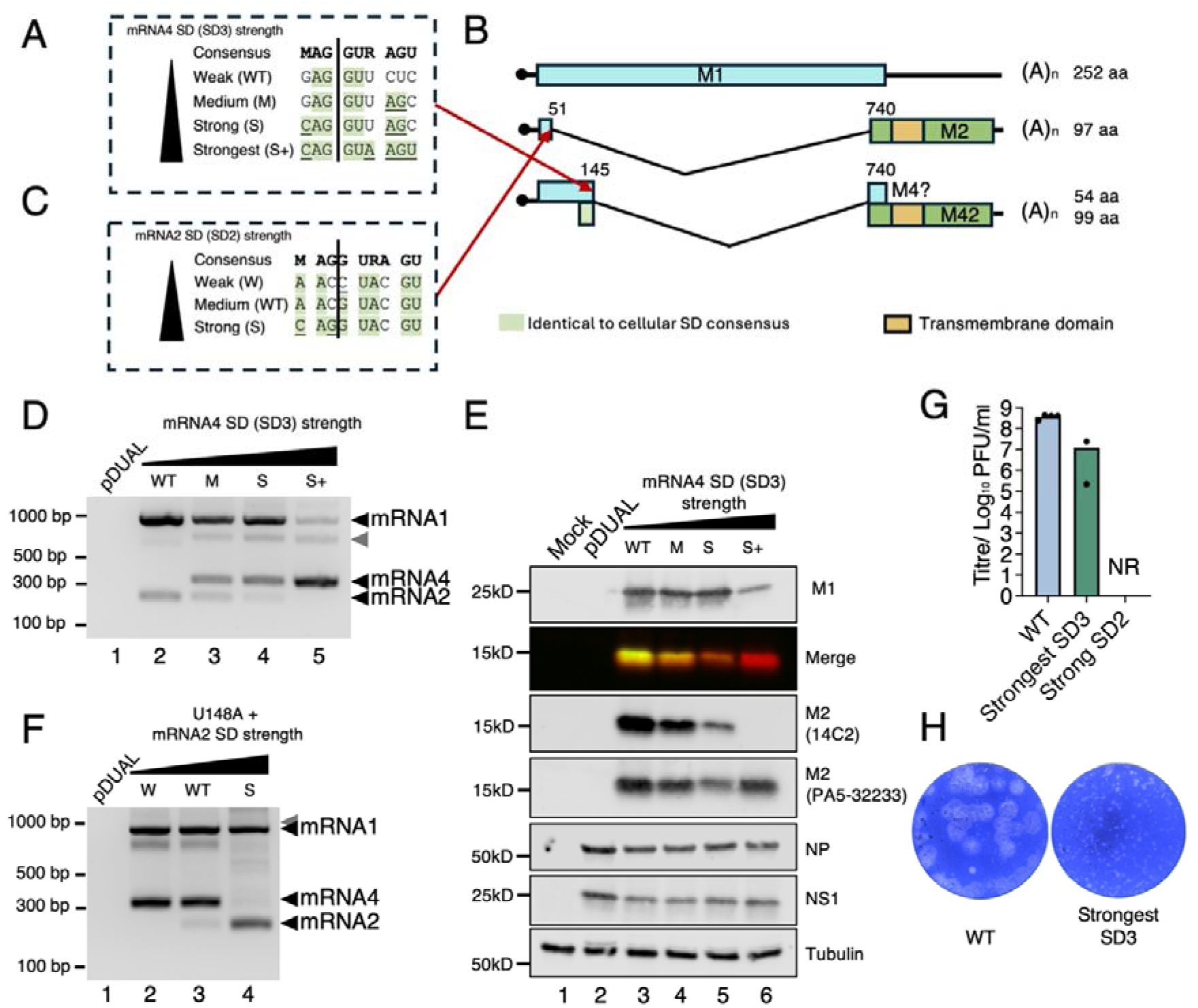
Splice site migration in segment 7 does not necessarily require inactivating mutations in the original SD motif. (A-C) Design of mutations (underlined) to alter recognition of SD2/ SD3 by cellular spliceosome component U1 snRNA. Nucleotides identical to the cellular SD consensus are shaded in green, and mutations are underlined. (D-F) Mini-replicon assay of WT and mutant segment 7 gene expression. HEK 293T cells were transfected with IAV mini-replicon components, NS segment and the indicated segment 7 or pDUAL empty vector plasmids. Total RNA was extracted from cells 48 hours after transfection, and segment 7 mRNA was detected using RT-PCR followed by agarose gel electrophoresis (D and F). The grey arrowhead indicates a probable heteroduplex of mRNA1 and 4 PCR products. (67) Protein expression was analysed by SDS-PAGE and western blotting (E). (G) P1 stock titres of WT and mutant viruses. MDCK cells were infected with rescued viruses at an MOI of 0.001, and supernatants were harvested 48 hours post-infection (h p.i). Data are the mean ± SEM of at least two independent virus rescues. (H) Representative plaque assays (in MDCK cells) of WT PR8 and a mutant that relies only on mRNA4 production.

To test if the competition between mRNA2/SD2 and mRNA4/SD3 was reciprocal, we next examined mutations that either strengthened or weakened SD2 (Figure 2C) in the context of the activated mRNA4 splicing induced by introducing an activating U148A substitution in mRNA4 SD3 (48). MaxEnt-predicted strengths ranged from 1.24 to 10.65, in comparison to the WT 9.51 (Supplementary Figure 1A). The RT-PCR results indicated mRNA4 SD activated by U148A blocked the weak and medium strength mRNA2 SD (Figure 2F, lane 2 and 3), while a strong SD2 outcompeted this SD3 (Figure 2F, lane 4). Thus competition between the SD2 and SD3 is reciprocal, supporting the possibility that a change in SD usage could be driven by gradual mutational changes in the intrinsic strength of cryptic and existing SD sites.

Although the mini-replicon system is a reliable model for studying IAV mRNA transcription and splicing without obstruction from M1 or M2 dysfunction arising from non-synonymous mutations, it does not reveal how changes in splicing patterns affect virus fitness. Therefore, to test whether the virus could tolerate the SD shift, as well as to explore potential evolutionary conflicts between the multiple functional elements in segment 7, we attempted to rescue the two mutants that utilised full-strength SDs to outcompete the other SD. Viruses were generated by transfecting bidirectional reverse genetics plasmids into HEK 293T cells, any progeny amplified by one passage in MDCK cells, and then plaque titred. WT PR8 grew normally to over 10^8^ PFU/ml and produced large plaques (Figures 2G, H). A virus with the strongest SD2 sequence could not be rescued, whereas the strong SD3 mutant could, but gave titres at least 10-fold less than that of WT and produced pinpoint-sized plaques indicative of impaired viral fitness.

Overall, the mRNA2 SD (SD2) and mRNA4 SD (SD3) could be “evolved” to outcompete each other through increased strength without additional inactivating mutations. Notably, only two nucleotide substitutions (causing two amino acid changes in M1) were sufficient to allow SD3 to outcompete SD2 and produce a viable virus, whereas three to five nucleotide substitutions (also causing two amino acid changes) were required to completely switch SD usage to SD3, but these were not compatible with virus rescue.

### 3’-proximal SD sites do not outcompete the mRNA2 SD

Both SD2 and SD3 sites are near the 5’-end of the M1 coding region, so although it was possible to swap their usage, this did not recapitulate the ICV/IDV strategy of an SD near the 3’-end of the M1 coding sequence and a consequently short intron. However, two avian IAV strains have been reported to use (at low levels) SD4 to produce mRNA5 (56), while our own bioinformatic analysis predicted a potential SD5, both located closer to the 3’-end of M1 and thus leading to relatively small introns (Figure 1B). Shorter introns are proposed to have a higher splicing efficiency (68), so to test whether these more 3’-proximal SD could also outcompete SD2, we used the same strategy of mutationally improving the match of the SD sequences to U1 snRNA.

For SD4/mRNA5, we introduced 5 nucleotide changes, leading to two amino acid changes in M1, that made the sequence a perfect match to cellular consensus and increased its MaxEnt score from a weak value of 0.78 in the WT segment to 10.9 (Figure 3A; ‘Strong SD4 mutant, Supplementary Figure 1B). Correspondingly, testing construct activity by RT-PCR in the mini-replicon system detected a new spliced transcript, mRNA5 from the mutant segment (Figure 3B, lane 3). Sequencing of the RT-PCR product confirmed use of SD4 instead of SD2 (Supplementary Figure 2A, B). However, unlike when SD3 was improved to perfectly match the cellular consensus for binding to U1 snRNP, SD4 did not outcompete the SD2. Consistent with this, western blotting showed apparently normal M2 synthesis, although a novel faster-migrating version of M1 (“M5”) that reacted with antisera raised against full length M1 but not serum raised against its C-terminal region was identified (Figure 3C lane 4). Therefore, to mimic a two-step migration event in which the gain of an SD is followed by loss of the original SD, we combined the activating mutations for SD4 with the G52C inactivating mutation for the old SD. When mRNA2 production was downregulated due to its impaired SD, increased mRNA5 and M5 expression was seen (Figure 3B lane 4), as well as a drastic reduction of M2 expression (Figure 3C, lane 5), indicating successful swapping of SD sites. We next investigated whether the virus could tolerate the splice site migration from the dominant SD (mRNA2 SD) to the weaker 3’-proximal SD of mRNA5 (SD4). However, a virus with the strong SD4 did not generate a viable virus (Figure 3D), perhaps at least in part because it carried two non-synonymous mutations in M1 (in M165Q and T167S; Figure 3A). Therefore, we designed a further SD4 mutant which only carried the T176S mutation but which still gave a high MaxEnt score for SD4 (Supplementary Figure 1B). When tested in the mini-replicon system, it produced readily detectable amounts of mRNA5, as well as slightly reduced amounts of mRNA2 (Figure 3E, sequencing results in Supplementary Figure 2C).

**Figure 3.**
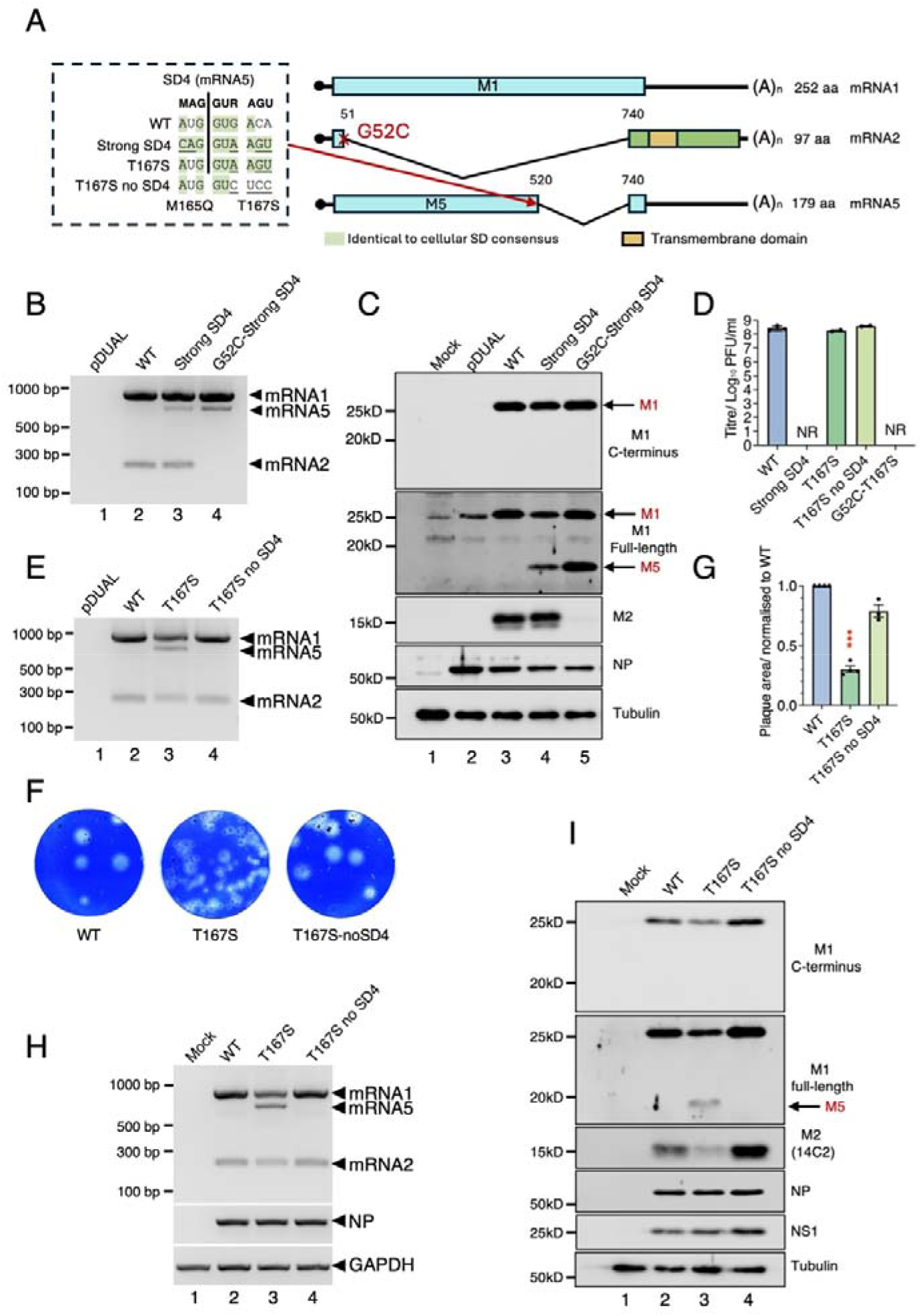
A 3’-proximal SD site does not outcompete the mRNA2 SD. (A) Mutational strategy for activating an alternative 3’-proximal SD, mRNA5/SD4 in PR8 segment 7. Identical nucleotides to the cellular SD consensus are shaded in green. Mutations are underlined. (B, C, E) HEK 293T cells were co-transfected (or mock transfected) with IAV mini-replicon components, NS segment and the indicated segment 7 or pDUAL empty vector plasmids, harvested at 48h and analysed by (B, E) RT-PCR and agarose gel electrophoresis and (C) western blotting. The novel M5 protein translated from mRNA5 is shown by an arrow. (D) End-point titres at 48h p.i. of PR8 WT and mutant viruses grown in MDCK cells (MOI = 0.01). Data are the mean of at least two independent rescues. NR = not rescued. (F) Representative images of plaque assays of WT and indicated mutants. (G) Plaque size quantification of PR8 WT and mutant viruses. Data are the mean ± SEM of 5-20 plaques normalised to the average WT value from at least three independent experiments. (H, I) The effects of SD4 mutations on viral mRNA splicing. MDCK cells were infected (or mock-transfected) with the indicated viruses at an MOI of 5. (H) Total RNA was isolated at 6 h p.i and the indicated mRNAs were detected by RT-PCR or (I) cells were lysed at 8 h p.i. and viral protein expression analysed by SDS-PAGE and western blotting.

This mutant was successfully rescued and grew to high titres but formed somewhat smaller plaques than WT virus (Figures 3D, F and G), suggesting that it tolerated the additional splice site well. However, once the mRNA2 SD/SD2 inactivating mutation G52C was introduced, the mutant was apparently non-viable (Figure 3D), presumably because mRNA5 only expressed M5 and not an ion channel protein (Figure 1B). To distinguish the impact of mRNA5 splicing from the M1 amino acid substitution on virus fitness, we generated a T167S derivative with an SD4 predicted to be non-functional (T167S no SD4; Figure 3A). In a mini-replicon assay, this construct did not produce an additional mRNA5 transcript, unlike its T167S parent (Figure 3E). Following rescue, the T167S no SD4 virus exhibited WT plaque size and titres (Figure 3D, F and G), indicating that attenuation was primarily associated with mRNA5 splicing rather than the M1 substitution. Infection of MDCK cells showed reduced mRNA1/M1 and mRNA2/M2 levels in T167S virus compared with WT (Figure 3H and I), suggesting that the fitness penalty might arise from the activation of an additional SD perturbing the normal M1/M2 splicing balance. Thus although only three nucleotide changes and one amino acid substitution in M1 were needed activate mRNA5 splicing, SD4 did not outcompete the dominant SD2.

To test another potential SD located towards the 3’-end of the M1 ORF whose use would lead to a relatively small intron, we turned to SD5 (Figure 1B). Unlike SD4, SD5 is already a good match to the cellular consensus and gave a strong MaxEnt score (Supplementary Figure 1C). However, it does not appear to be efficiently used, as no distinct band migrating at the expected size was detected in WT PR8 (Supplementary Figure 3B, lane 2). However, a novel spliced mRNA6 transcript was readily detectable after 2 nucleotide changes were added to SD5 to create a perfect match to the cellular consensus, using RT-PCR (‘Strong SD5’; Supplementary Figure 3B, lane 3, sequencing result in Supplementary Figure 2D). Although mRNA6 was predicted to encode a 16.8 kDa truncated M1, no such translation product was detected by western blotting (Supplementary Figure 3C lane 4). However, slightly altered migration and strongly reduced accumulation of M1 as well as a slight reduction in M2 expression was observed in this mutant. Furthermore, the Strong SD5 mutant could not be rescued on any of 5 independent attempts (Supplementary Figure 3D), indicating a severe fitness penalty. The Strong SD5 mutant contains two non-synonymous mutations in M1: L146Q and C148S. To disentangle which mutation led to impaired M1 and/or M2 accumulation, the two single mutants were generated and tested for their mRNA and protein expression. Neither single mutant produced detectable mRNA6 (Supplementary Figure 3B, lanes 4 and 5), but a similar reduction in M1 and M2 levels to the double mutant was seen in the L146Q mutant (Supplementary Figure 3C lanes 4 and 5), while C148S showed normal M1 and M2 (lanes 3 and 6). Thus, the impaired M1 and M2 accumulation was attributable to the L146Q substitution in M1, rather than the C148S mutation or mRNA6 production. Thus, like SD4, SD5 was unable to outcompete the canonical SD2 and furthermore, its activation to produce detectable amounts of mRNA6 incurred a high mutational cost on virus fitness because of a nonsynonymous change in M1. Together, our results in this section suggest that SD relocation is possible but constrained. Even though a functional alternative SD can be activated, multiple nucleotide changes are required and complete replacement of the canonical SD is dependent on the wider context of the new SD. Further evolutionary progression is limited by constraints imposed by multiple factors influencing viral fitness, including but not limited to, M1 and/or M2 coding sequences.

### Examining splice acceptor site migration during influenza virus evolution

So far, we investigated the evolutionary flexibility of SD usage in the IAV matrix segment. The next question to address was around the potential mobility of the segment SA site. A complete SA motif involves three separate functional units whose spacing and order matter: from 5’- to 3’, a BP motif, a PPT and then the SA site itself (36, 38, 39). Minimally, this involves a total of around 30 ntds, in contrast to a maximum SD site of 9 nucleotides, potentially making an SA unit more difficult to alter. IAV segment 7 sequences, in the M2 coding region (to match the approximate location of the SA motif in ICV and IDV), were inspected to find potential cryptic or relic SAs. This highlighted a potential PPT located between ntds 839 and 867 along with a potential adenosine branch point residue (Supplementary Figure 4A). To investigate whether this could form a functional SA unit, two mutations C870A and U872G, synonymous in M2, were introduced to form a potential SA 133 nucleotides downstream of the canonical SA (Supplementary Figure 4A, predicted splice site strength in Supplementary Figure 1D). In the mini-replicon assay, mRNA2 levels in the C870A-U872G mutant reduced markedly and a new spliced transcript (named mRNA7) was apparent (Supplementary Figure 4B lane 3). Sequencing of the PCR product confirmed that the new group of 3’ splice elements had been successfully activated (Supplementary Figure 2E) and, concomitantly downregulated usage of the canonical SA. Furthermore, the C870A-U872G mutant virus could be successfully rescued, but it exhibited reduced fitness, characterised by a 10-fold reduction in viral titre and a small-plaque phenotype (Supplementary Figure 4C-E). When segment 7 mRNA splicing and protein expression were retested in infected MDCK cells, NP mRNA levels in C870A-U872G-infected cells were slightly lower than that of cells infected by WT (despite equal MOI), suggesting the mutant virus could not infect cells as efficiently as WT. RT-PCR analysis indicated a marked reduction in mRNA2 and the gain of mRNA7 (Supplementary Figure 4F lane 3), consistent with the mini-replicon data. Western blotted confirmed reductions in NP and M1 expression, as well as the reduction of M2 expression below the limit of detection (Supplementary Figure 4G). Together, these data showed that only two synonymous mutations could activate a latent 3’ splice unit, supporting the premise that SA sites can move during virus evolution.

The cryptic SA described above resulted in a spliced mRNA predicted to encode and M2 variant lacking a TM domain and therefore unlikely to function as an ion channel. We therefore searched for other cryptic SAs that would preserve more of the M2 coding region. This showed a CAG motif 9 nucleotides downstream of the active SA as a candidate with the potential for reactivation (Figure 4A). Usage of the downstream CAG is difficult to detect by PCR as it would only lead to a 9 ntd difference in fragment size, but sequencing of the WT segment 7 mRNA2 did not provide evidence of its usage, despite the downstream CAG also matching the SA consensus (Supplementary Figure 2A). Thus, to test whether the downstream SA was inhibited by the canonical SA, the latter was inactivated by the addition of an A738G change (Figure 4A) and the mRNA splicing patterns analysed by mini-replicon, and RT-PCR. This showed no detectable spliced mRNA products (Figure 4B lane 4), whereas WT produced the expected mix of mRNA1 and mRNA2, indicating the downstream cryptic SA remained inactivate. To enhance the intrinsic strength of the cryptic SA, we introduced a further A749G substitution, in the context of the A738G mutation. However, no splicing was detected in the A738G-A749G double mutant (Figure 4B lane 6), indicating that a stronger cryptic SA alone was still insufficient. However, the downstream CAG is a further 9 ntd away from an already sub-optimal PPT, so we therefore introduced additional pyrimidine substitutions in the PPT upstream of both canonical and cryptic SAs (labelled as “pY”), to enhance splicing efficiency (42, 69). In an otherwise WT segment 7, the polypyrimidine substitutions modestly increased usage of the canonical SA (Figure 4B lane 3). However, when the canonical SA was inactivated by A738G and the PPT was strengthened, splicing was now detected from the downstream CAG site (pY + A738G; Figure 4B lane 5 and Supplementary Figure 2F). Splicing from the downstream CAG sequence was further increased when the cryptic SA was additionally strengthened by the A749G substitution (Figure 4B lane 7 and Supplementary Figure 2F). When segment 7 splicing was moved from the canonical to downstream cryptic SA, the novel spliced transcript was predicted to encode an M2 variant with a shorter ectodomain, which would be unlikely to react with the ectodomain-specific monoclonal antibody 14C2 (Figure 4C). Protein expression from the alternative SA constructs was therefore analysed by western blotting using the N- and C-terminal domain-specific anti-M2 antibodies. As expected, the pY mutant gave a higher M2 expression level than WT, confirmed by reactivity with both anti-M2 antibodies, while the splicing null mutants A738G and A738G-A749G gave no detectable reactivity with either (Figure 4D, lanes 4, 5 and 7 respectively). Although the alternatively spliced transcript from the pY+A738G segment could be detected, any M2 variant expression was below the detection limits of both antibodies, perhaps due to the low splicing efficiency and/or lowered M2 stability (lane 6). However, an M2 polypeptide from the pY+A738G-A749G construct was detected by the polyclonal anti-M2 serum but not the 14C2 antibody, consistent with it possessing a shorter ectodomain resulting from usage of the downstream SA (lane 8). Overall, these results demonstrate that the downstream cryptic SA can be reactivated by strengthening the SA unit while inactivating the canonical SA, revealing latent plasticity in segment 7 SA usage.

**Figure 4.**
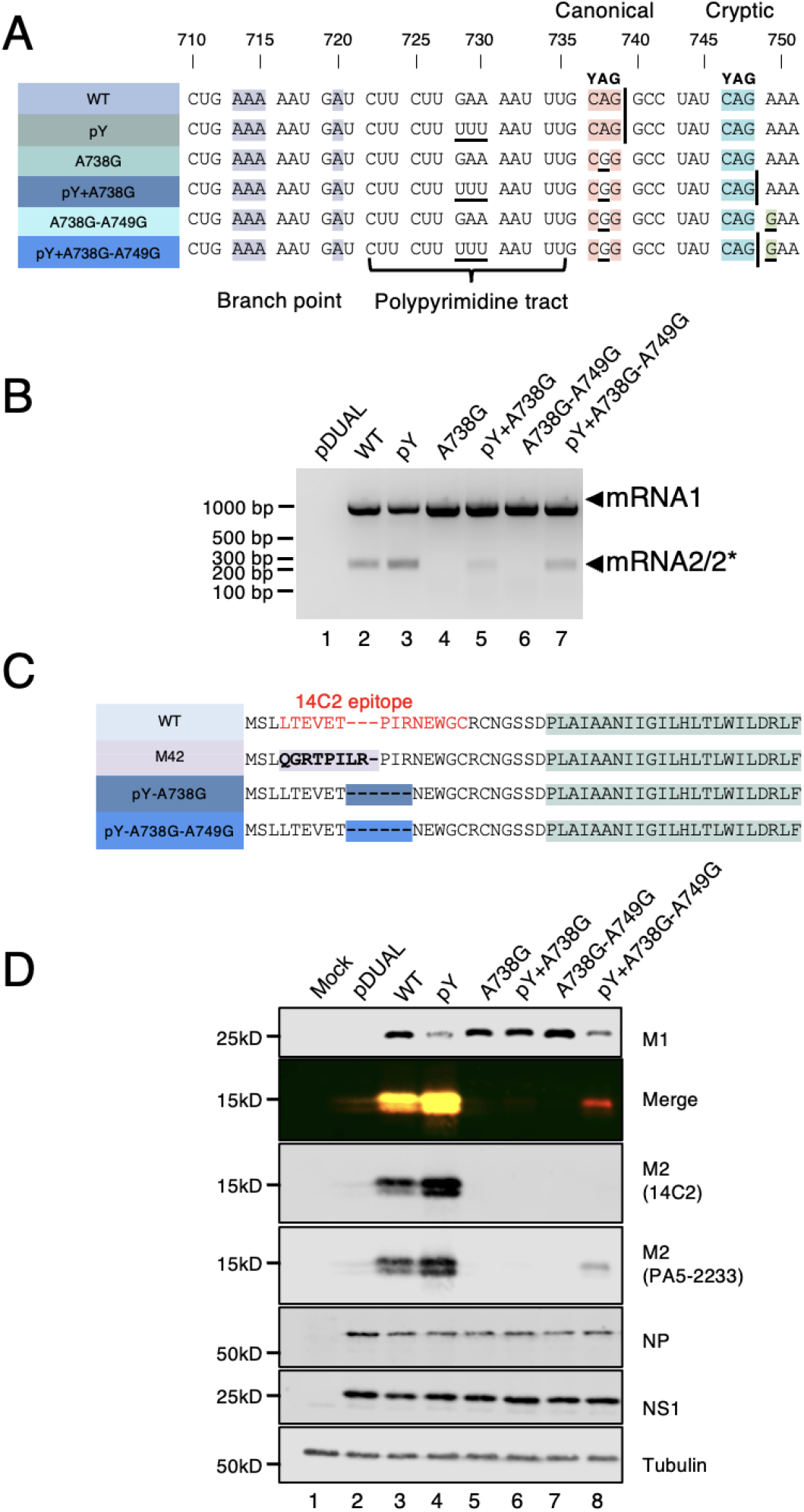
Activation of a downstream cryptic SA in segment 7. (A) Mutational strategy for activating a downstream cryptic SA in segment 7. The BP is highlighted in purple, the canonical SA in orange and the cryptic SA in blue. The mutation A749G, predicted to increase the strength of the cryptic SA (42) is highlighted in green. Vertical lines mark exon-intron boundaries. Mutations are underlined and the segment 7 PPT is labelled. (B and D) HEK 293T cells were co-transfected (or mock transfected) with the mini-replicon components, NS segment and the indicated segment 7 or pDUAL empty vector plasmids, harvested at 48h and analysed by (B) RT-PCR and agarose gel electrophoresis and (D) western blotting as labelled. * indicates co-migrating bands corresponding to the spliced products generated from the cryptic SA. (C) Alignment of the predicted N-terminal sequences of M2 and variants resulting from downstream SA activation. Residues included in the 14C2 antibody epitope are indicated in red (48, 70). The transmembrane domain of M2 is shaded in green.

To determine whether use of the downstream cryptic SA was compatible with viral replication, the viability of mutants designed to activate it was tested by virus rescue. The A738G and A738G-A749G mutants which failed to express detectable mRNA2 and M2 in the mini-replicon system were non-viable. In contrast, the pY+A738G and pY+A738G-A749G mutants that utilised the cryptic SA were viable, but gave lower titres and exhibited smaller plaques compared with WT (Supplementary Figures 5A-C). Next, the viral mRNA splicing and protein expression were analysed by single-cycle infection. In cells infected by WT PR8, the expected mixture of mRNA1 and mRNA2 was detected (Supplementary Figure 5D lane 2). In contrast, cells infected with pY-A738G-A749G produced mRNA1 together with an additional slightly faster-migrating RT-PCR product (lane 4). Sequencing confirmed that this transcript resulted from splicing at the downstream cryptic SA (Supplementary Figure 2F), consistent with the mini-replicon results. Western blotting showed that cells infected with pY-A738G-A739G expressed an M2-related polypeptide that was detected by PA5-32233 but not by 14C2 (Supplementary Figure 5E lane 4). Therefore the downstream cryptic SA could be activated by 5 ntd changes in the context of virus infection to express an M2-variant with an altered ectodomain and 3 amino-acid changes in M1, further supporting the possibility of SA site migration during influenza virus evolution.

### Moving SD and SA sites simultaneously to produce a new transcript that encodes an ion channel

The results above show that IAV can tolerate changes to alternative SD or SA usages made individually. However, our hypothesis is that during evolution of the orthomyxovirus matrix segment splicing pattern, both SD and SA sites have moved. Therefore, we next examined whether IAV could tolerate simultaneous changes in both SD and SA sites. Among the potential SD candidates, usage of SD1/mRNA3 would result in the loss of all start codons whose use would initiate translation of the M2 ORF upstream of the TM domain. The mutations needed to activate SD4/mRNA6 SD were not tolerated by the virus. Therefore, SD3/mRNA4 and SD5/mRNA5 were selected for further analysis. Regarding the choice of new SA, it should locate upstream of approximately nucleotide 786 of segment 7, which encodes the N-terminal boundary of the M2 transmembrane domain, essential for ion channel function (14, 71).

Therefore, to establish an ICV/IDV-like splicing strategy in IAV, our first attempt was to activate the mRNA5 SD (using the ‘Strong SD4’ mutations described in Figure 3A) and move the SA upstream of the canonical SA to place the transmembrane domain coding region in frame with the M1 ORF. CAG triplets were introduced upstream of the natural segment 7 SA to generate novel SAs which would place the main M2 exon in frame with the M1 ORF, which would produce novel spliced mRNAs predicted to encode first 165 amino acids of M1 fused (via a variable linker sequence) to amino acids 9-97 of M2; an ‘M52’ polypeptide (Supplementary Figure 6A). These constructs were then tested in the mini-replicon system.

Of the three candidate SA sites tested, SA1 and SA2 were nonfunctional, whereas SA3 produced an mRNA5-size and an mRNA2-size transcript (Supplementary Figure 6B, sequencing result in Supplementary Figure 2G). Viral protein analysis showed no normally sized M2 polypeptide was detected by either the 14C2 or C-terminal M2 antibodies (Supplementary Figure 6C), suggesting the new SA blocked the natural one and moved SD2 out of frame with the M2 ORF. However, a ∼28 kDa polypeptide translated from mRNA5 was detected by anti-M2 polyclonal antibody PA5-32233 (Supplementary Figure 6C lane 7), representing the M52 fusion protein. When virus rescue was attempted, neither the Strong SD4-SA3 construct or the T167S-SA3 variant (with a slightly weaker SD4 but only a single nonsynonymous change in M1; see Figure 3) resulted in viable virus (Supplementary Figure 3D). Thus, although the re-wired segment 7 could generate an M5-M2 fusion protein with some similarities to the P42 protein produced by ICV and IDV, it could not functionally replace the native IAV M2 expression strategy.

A reason why a segment 7 that only expressed M52 was non-viable might be the failure of the polyprotein to insert into the membrane, because of the oversize ectodomain and/or the lack of a signal peptide to direct its cleavage. Adding an SP motif would increase the number of mutations required to move expression from the IAV to the ICV/IDV pattern, further reducing the likelihood that it could occur via natural evolution. Therefore, we reverted to testing novel splicing strategies in which the more 5’-proximal SD3 replaced SD2, as depending on the exact strategy, this would either lead to a smaller ectodomain or the use of the M42 initiation codon. To achieve this switch, two synonymous substitutions in M1, G52C and U148A, were introduced to inactivate the mRNA2 SD2 and promote usage of the mRNA4 SD3, respectively (42, 48). In this background, three distinct strategies were designed to alter the SA site (Figure 5A). Previous studies have shown that CAG triplets located a short distance upstream of the canonical SA can function as an alternative SA (42, 72). Therefore, in strategy (i), the CAG triplets (SA1-SA3) were introduced within an 8-nucleotide window upstream of the canonical SA such that splicing at them SAs would place the M2 coding region in frame with the M1 ORF. Translation initiation at the M1 AUG codon would produce an M1-ion channel fusion protein, analogous to the ICV/ IDV strategy (Figure 5A (i)). In the second strategy, three SAs (SA4-SA6) were introduced into the same window upstream of the canonical SA, but so that any spliced transcripts were in frame with the M42 AUG codon, so that translation initiated via leaky ribosomal scanning using AUG at 114-116 nucleotide of segment 7 would produce an ion channel with a variant ectodomain (Figure 5A(ii)). The third strategy aimed to produce another M42-variant peptide through activation of the downstream cryptic SA (SA7). This was achieved by inactivating the canonical SA, enhancing the PPT and strengthening the context of SA7 (Figure 5A(iii)). The alternative splicing site activation and resulting splicing patterns were first examined using transfection-based mini-replicon assays and RT-PCR. The mutant G52C-U148A was included as a positive control (48) and the detection of mRNA4 confirmed successful SD3 activation (Figure 5B, C; ‘Natural SA’, Figure 6D, lane 3). On the background of SD3 activation, novel spliced transcripts were detected from the all the SAs except SA1 (Figures 5B-D). Compared to mRNA4 (Figure 5, SD3-Natural SA), slightly slower migration rates were observed for the SA2-SA6 spliced transcripts, consistent with the usage of SD3 in combination with the alternative SAs. Sequencing analysis validated that these spliced transcripts did indeed use the SD3 in combination with the novel SAs (Supplementary Figure 2H). The spliced transcripts were predicted to encode M1-M2 fusion proteins (Figure 5E) or M42-variants (Figure 5F) which contained a transmembrane domain and variable ectodomains resulting from the alternative SD and SA utilisation. We next examined whether these spliced transcripts were translated into predicted viral proteins by western blotting and double M2 antibody staining as before, expecting that the novel viral proteins would only react with PA5-32233. M2 protein expressed by WT segment 7 reacted with both anti-M2 antibodies (Figure 5G, lane 3). The mutant G52C-U148A, which expressed M42 only, was used as a positive control for a protein that only reacted with the C-terminal M2 antibody (lane 7). In mutants spliced using the mRNA4 SD and SA2 or SA3, ∼15 kDa peptides were detected by PA5-32233 but not 14C2 (lanes 5 and 6). The apparent sizes of the novel proteins were larger than WT M2 and M42, consistent with their larger 2^nd^ exons due to the splice site changes. In addition, the SA3 mutant exhibited stronger protein expression than SA2, in agreement with its higher splicing efficiency observed by RT-PCR. Although weak splicing was observed in the SA4 construct, M2 expression did not reach the antibody detection limit (Figure 5H, lane 4), but SA5 and SA6 western blotting with PA5-32233 showed a good correlation of ion channel expression and spliced mRNA abundance (lanes 5 and 6). Loss of reactivity with 14C2 confirmed that these segment 7 mutants produced M42 variants with distinct ectodomains compared to WT M2. For the new SA downstream of the canonical SA (SA7), the two mutants only containing either an activated SD3 or SA7 (pY-A738G-A749G), were used as controls to confirm the novel splice site activation and resulting change in the M2 ectodomain (Figure 5F) by lost reactivity with the 14C2 antibody but not PA5-32233 (Figure 5I lanes 4 and 5). Similarly, the mutant using both novel SD3 and SA7 showed reactivity with PA5-32233 rather than 14C2 (Figure 5I lane 6). This result confirmed that a novel viral protein with an identical C-terminal domain to M2 but a distinct ectodomain was expressed, consistent with the RT-PCR and sequencing data.

**Figure 5.**
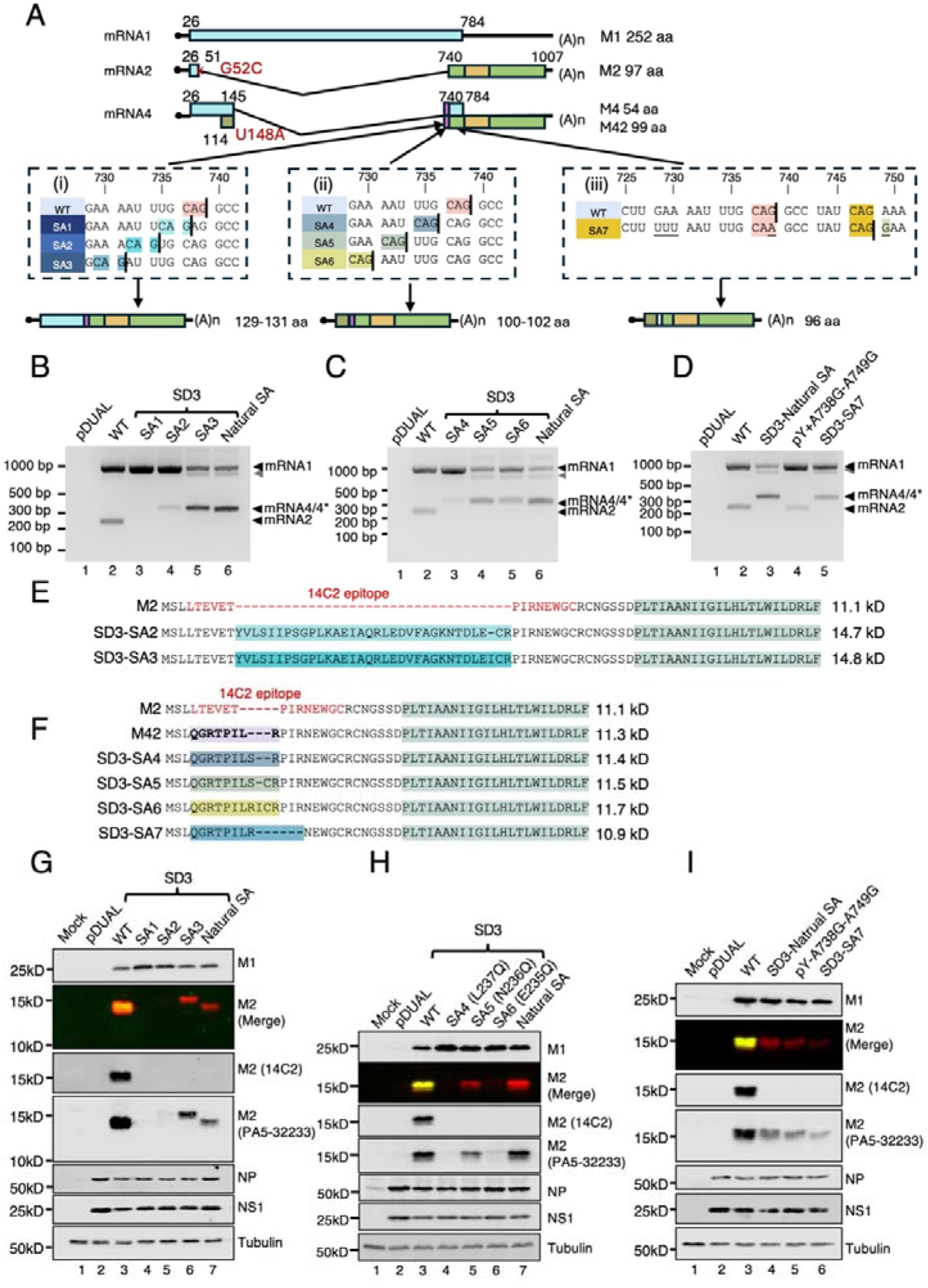
Cryptic SD and SA sites can be simultaneously activated to produce novel ion channel proteins. (A) Diagrammatic summary of mRNA splice variants. The nucleotide coordinates of ORFs, SD, canonical and upstream cryptic SA sites are labelled. Potential ORFs are colour coded (blue, M1; light green, M2, deep green, M42 N-terminus) and the transmembrane domain in M2 is highlighted in yellow. The additional region produced by SA alteration is highlighted in purple. The deleted exon is indicated by a white box. Dashed boxes give key sequences in triplets corresponding to M1 ORF. The canonical splice acceptors are highlighted in orange, while novel CAGs are highlighted in different shades of blue, grey and yellow. Schematics (i), (ii) and (iii) illustrate strategies 1, 2 and 3, respectively. (B-D, G-I) The mini-replicon components, NS segment and the indicated segment 7 or pDUAL empty vector plasmids were co-transfected (or mock-transfected) with the indicated segment 7 mutants in HEK 293T cells and (B-D) segment 7 mRNA detected by RT-PCR followed by agarose gel electrophoresis or (G-I) viral protein expression analysed using SDS-PAGE and western blotting. * indicates co-migrating bands corresponding to spliced products generated from alternative SAs. The grey arrowhead indicates a probable heteroduplex of mRNA1 and 4/4* PCR products. (E, F) Alignment of the predicted N-terminal sequences of M2 and variants resulting from rewired splicing patterns. The range of residues implicated in recognition of M2 by the 14C2 antibody is indicated in red (48, 70). The transmembrane domain of M2 is shaded in green. (G-I).

**Figure 6.**
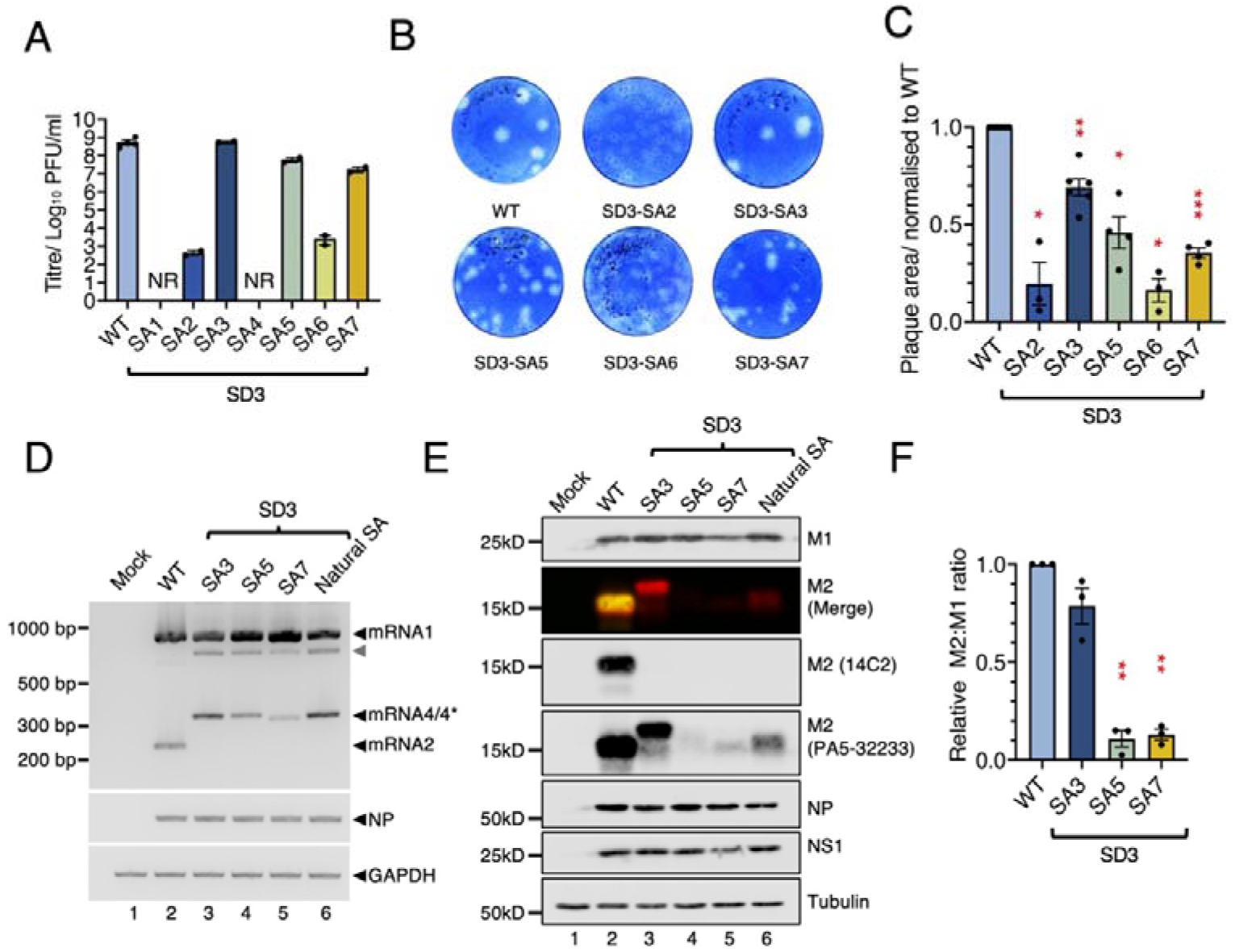
Viruses with using alternative segment 7 SD and SA sites are viable. (A) 293T cells were transfected with reverse genetic plasmids to create the indicated viruses and the supernatant passaged once in MDCK cells to generate P1 stocks. Data are the resulting titres and represent the mean of at least two independent rescues. NR = not rescued. (B) Representative images of plaque assays of WT and indicated mutants. (C) Quantification of the plaque size of PR8 WT and mutant viruses. Data are the mean ± SEM of 5-20 plaques normalised to the average WT value from at least three independent experiments, compared using an ordinary One-way ANOVA test with Dunnett’s post-test (N=4-6). **** = P<0.0001; *** = P<0.001; ** = P<0.01; * = P<0.05; absence of significance symbols = P>0.05. (D and E) MDCK cells were infected (or mock-infected) with the indicated viruses at an MOI of 10 and (D) total RNA isolated at 8 h p.i and the indicated mRNAs detected by RT-PCR, or (E) cells were lysed at 16 h p.i and viral protein expression analysed by SDS-PAGE and western blotting. The grey arrowhead indicates a probable heteroduplex of mRNA1 and 4/4* PCR products. * indicates co-migrating bands corresponding to the spliced products generated from alternative SAs. (F) M2 isoform:M1 ratios were quantified by densitometry of PA5-32233 reactivity from replicate experiments and normalized to the value of WT construct. The relative M2: M1 ratios were analysed by an ordinary One-way ANOVA test with Dunnett’s post test relative to WT (N=3-7). **** = P<0.0001; *** = P<0.001; ** = P<0.01; * = P<0.05; absence of significance symbols = P>0.05. Error bars indicate the standard deviation.

Using SD3 and a variety of novel SA sites led to the successful production of multiple new spliced transcripts which encoded a variety of putative ion channels in the mini-replicon system. The question therefore arose as to whether the virus could tolerate the new splicing patterns. Rescue of the entire panel of mutants was therefore attempted and any progeny amplified by one passage in MDCK cells. WT PR8 grew to an average titre of (PFU/ml (Figure 6A). Most mutants were successfully rescued, except SA1 and SA4, perhaps reflecting their lower splicing efficiency and subsequent ion channel protein shortage. In addition, the non-synonymous mutations which were introduced for cryptic SA activation in M1 might negatively impact the virus’s fitness (Supplementary Table 1). Of the mutants which successfully rescued, SA2 and SA6 exhibited severe replication defects, with titres much lower than that of WT. SA3 gave WT-level titres, whereas SA5 and SA7 replicated approximately 10-fold lower than WT. In addition, small-plaque phenotypes were observed in all mutants (Figure 6B-C). Thus, the virus could tolerate altered splicing patterns, although the mutants exhibited varying levels of fitness defects. To provide biochemical evidence for the alternative splicing patterns, we next examined segment 7 mRNA splicing by the viruses which grew to sufficient titre to allow high multiplicity infection of MDCK cells. Following infection, total RNA was extracted and analysed using RT-PCR. The abundance of GAPDH mRNA amplicons indicated equal recovery of material (Figure 6D). Levels of NP mRNA were also comparable between infected samples, suggesting that all infections had proceeded successfully. However, clear differences in segment 7 splicing patterns were seen. In the cells infected by WT virus, both unspliced mRNA1 and spliced mRNA2 were present (Figure 6D lane 2), while no mRNA2 was detected in any of the mutants with an SD3 background due to the loss of mRNA2 SD site via the G52C mutation. The SD3 mutant using the natural SA site generated unspliced mRNA1 and spliced mRNA4 rather than mRNA2, as expected (lane 6). The three mutants with novel SA sites also generated mRNA4-sized products although there was a slight increase in migration rate for the SA7 construct, consistent with its larger intron. Sequencing of the various PCR products confirmed use of the variant splice sites (Supplementary Figure 2I). To further assess the impact of alternative splicing on M segment-encoded protein expression, viral proteins were analysed using western blotting (Figure 6E, quantitation in 6F). Cells infected with the WT virus isolated contained a polypeptide recognised by the M2 ectodomain-specific 14C2 antibody, but as before, the cells infected with mutants using SD3 instead of SD2 did not; however, SA3, SA5, SA7 and SD3-natural SA produced detectable amounts of PA5-32233-reactive proteins (Figure 6E lanes 3-6), indicating changes to the M2 ectodomain. The M42 variant expressed by SA3 also showed an obvious size shift, as expected from the splicing pattern changes (Figure 6E lane 3). Expression of all the M42 variants was lower than that of M2 (as expected for products made by leaky ribosomal scanning) but varied significantly (relative to M1 levels) between constructs (Figure 6F). SA3 apparently expressed better than the parental M42, whereas SA5 was near the detection limit of PA5-32233 antibody. As the levels did not correlate well with transcript abundance, differential protein stability might provide an explanation. Nevertheless, our data show that it is possible to rearrange segment 7 splicing patterns to generate novel forms of the ion channel whilst retaining virus viability, indicating a surprising degree of evolutionary flexibility.

## Discussion

We initiated the study reported here because we were interested in how the diversity of splicing patterns seen in influenza virus matrix segments might have evolved. We assume that the ancestral orthomyxovirus did not use mRNA splicing, as nuclear replication is a derived characteristic that sets orthomyxoviruses apart from most other negative sense RNA virus families. The last common ancestor (LCA) orthomyxovirus matrix segment need not have been functionally monocistronic; there are other means of expressing two polypeptides from one mRNA in eukaryotic cells than splicing. Influenza B virus for instance, uses a translation termination/reinitiation mechanism (31). However, the LCA of the current influenza viruses must at least have had an ORF located towards the 3’-end of the segment (in positive sense terms) that when expressed even at low level was beneficial and could be positively selected for (73). To reach the current gene structures seen in influenza viruses, this ancestral ORF minimally had to encode (or gain) a transmembrane domain that oligomerizes into an ion channel and an ectodomain that allows its membrane insertion, as well as a reasonably efficient expression mechanism. Our primary focus was on the expression mechanism for the three species that use splicing. In one possible model, splicing evolved once to create a matrix segment that efficiently expressed both M1 and M2, which then diversified to produce the subtly different ICV and IDV strategies and then further (by splice site migration) to the completely different pattern of IAV (Figure 7A). A second model is that the IAV and ICV/IDV splicing strategies evolved independently from an unspliced influenza virus matrix segment (Figure 7B).

**Figure 7.**
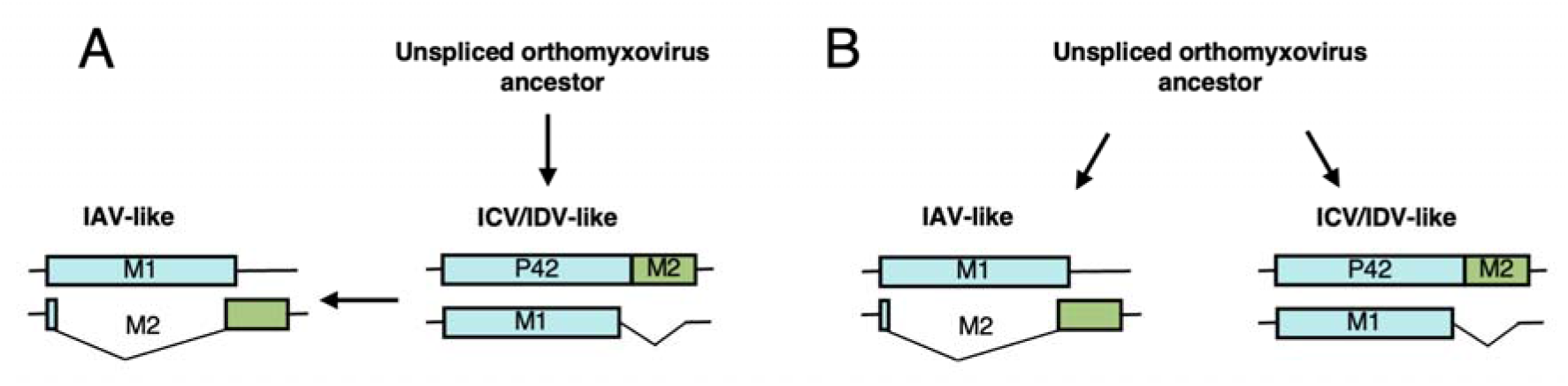
Two possible evolutionary pathways for matrix segment splicing strategies in influenza viruses. (A). An unspliced orthomyxovirus ancestor first evolved ICV/IDV-like splice strategies, followed by splice site migration to give an IAV-like strategy. (B). Independent evolutionary gains of splicing by unspliced ancestral orthomyxovirus(es) produced IAV- and ICV/IDV-like splicing strategies.

We acknowledge that our artificial system in which we ‘force evolved’ IAV using rationally designed reverse genetics constructs cannot recreate the actual evolutionary path followed by the matrix segment. However, it can provide insights into how easy or difficult (and therefore how likely) it is to gain and/or move splice sites in an orthomyxovirus segment. We found that IAV segment 7 exhibits substantial plasticity over its splicing patterns. We show that it contains at least five functional or cryptic SD sites that can be activated or inactivated by relatively small numbers of mutations, and that multiple SA sites can also be created. Furthermore, we show that viable viruses using these cryptic SD and/or SA sites can be produced, including ones that completely rewire the segment 7 splicing pattern. Overall, we interpret this as indicating that evolutionary gain of splice sites for a nuclear-replicating virus that piggy backs onto RNA polymerase II (25) is not intrinsically unlikely. This does not however distinguish between independent evolution of the IAV and ICV/D patterns, or diversification by migration of splice sites (Figure 7).

It is evolutionarily implausible for the splice site migration model (Figure 7A) to occur via simultaneous movement of both SD and SA sites, as too many mutations in multiple regions of the segment would be required. More plausible, is a stepwise shift involving the acquisition of alternative splice sites whose balance changes over time before settling into a new dominant pattern. We show here that PR8 IAV can tolerate the presence of at least two pairs of competing SD sequences (SD2 and SD3, or SD2 and SD4), consistent with a migration model. Competition between splice donor sites (SDs) in cellular mRNAs is known to be influenced by their intrinsic sequence strength (59, 74-76). SDs which possess enhanced complementarity to U1 snRNA have an increased likelihood of usage (74-77). Our data show that competition between IAV segment 7 SDs sometimes but not always follows a similar pattern. When SD3 was engineered to perfectly match the cellular consensus, it effectively outcompeted SD2, suggesting that intrinsic SD strength was the dominant determinant. However, neither SD4 or SD5 could completely displace SD2 even after being altered to a perfect consensus match, suggesting the involvement of additional splicing regulatory elements. This is consistent with several previous studies indicating that *cis*-acting regulatory elements can reinforce mRNA2 usage even in the presence of stronger alternative SDs (78). However, a role for additional splice site regulatory sequences does not rule out the splice site migration model, although it perhaps adds on an additional layer of evolutionary modification needed to produce a stable new pattern of splicing.

The segment 7s of all strains of IAV are thought to use at least two SD sites, but we are unaware of any strong evidence for alternative SA usage. Perhaps reflecting this, we saw a more binary choice between SA sites, with most novel SAs overriding usage of the canonical SA. For the series of novel SAs 1-6 added in or in between the PPT and canonical SA this was not unexpected, as in cellular mRNAs the first AG within the PPT region is generally used (72). The same logic explains why novel SA7 just downstream of the canonical SA only became active when the latter was mutationally inactivated. However, it is less clear why novel SA8 (C870A-U872G) further downstream of the canonical SA almost totally overrode normal 3’-splice site choice. Whether this results from the relatively weak PPT sequence upstream of the natural segment 7 SA which balances M1 and M2 expression (42) or from other context effects is unknown, but it might suggest that the migration model requires the gain of a novel SD site first.

Other major constraints on splice site evolution come from the overlapping coding regions. We have previously shown that most possible mutations in SD2 are not tolerated, probably because of non-synonymous changes in M1, though possibly also due to changes to the segment-specific packaging signal (42, 50). Here, as well as failing to rescue SD2 mutants again, we found that activation of SD5 was also incompatible with virus viability because of the concomitant L146Q change in M1. Conversely, many M1 mutations were tolerated. Overall, 11 separate mutants with a variety of non-synonymous changes in up to three codons were rescuable (Supplementary Table 1), albeit with varying degrees of fitness penalty. Thus although not all potential splicing events are compatible with viral viability, M1 coding constraints can be bypassed.

A further major constraint to splice site variation comes from the ion channel itself. We assumed that retention of the TM domain is a necessity, for ion channel formation and for the M2 cytoplasmic domain to be localized correctly to function in virus assembly (79). The finding that a virus using SA8 to the almost total exclusion of the canonical SA (a faint trace of mRNA 2 was visible from infected cells, although M2 protein was below the limit of detection) was not only viable but replicated to reasonable titres was therefore a surprise (Supplementary Figure 4 and Supplementary Table 2). Whether this virus is phenotypically near-normal due to the very low levels of residual M2 expression, and/or PR8 virus is less reliant on ion channel function than less laboratory adapted viruses remains to be determined. We note however, that in general, the expression levels of splice variant M2 channels did not correlate that well with virus titre (Supplementary Table 2). For the M2 variants that retained a TM domain, we assume that a further constraint arises from the fact that the M2 ectodomain influences membrane insertion (80). The exact sequence specificity of this process is not known except to note that it seems to be flexible as both the M42 and the paramyxovirus simian virus 5 small hydrophobic protein sequences also suffice to direct the protein to the membrane (48, 80, 81). Consistent with this, we found the M2 and M42 ectodomains to be broadly tolerant of insertions up to 34 amino acids, as well as small deletions (Supplementary Table 2). However, the strong SD4/ T167S-SA3 viruses with a much longer M1-derived ectodomain could not be rescued, perhaps reflecting that the TM domain needs to be either downstream of a signal peptide sequence or relatively near the N-terminus. Other than this however, the exact sequence of the M2 ectodomain does not seem to exert much constraint on splice site evolution. This is consistent with its unstructured nature and lack of strong evidence for any function other than membrane insertion (82).

Not only segment 7, but virus also exhibits tolerance to splicing pattern change of other segments. Previous studies have reported that the viral replication is not impacted by inserting an exogenous intron and splitting the ORFs of NS1 and NEP (83, 84). Apart from segment 8, normally non-spliced segments such as segment 5 and 6 can also harbour artificial introns and be rescued successfully (85). Moreover, the NA mutant exhibited high genome stability over four serial passages. These results demonstrate additional splicing events can potentially expand viral genome capacity. Beyond the switch from non-splicing to splicing, the virus also shows a degree of flexibility in spliced segment rearrangements. For example, the viable virus whose segment 7 and 8 were engineered to encode nonoverlapping independent M1/ M2 or NS1/ NEP ORFs, could be generated but exhibited reduced fitness (86). The tolerance and plasticity of splicing pattern switches in both spliced and unspliced segments reveal the evolutionary potential of splicing in influenza virus.

## Data availability

Primary data are archived on Figshare. Link available on reasonable request.

## Author Contributions

SZ: Conceptualisation, data curation, formal analysis, investigation, methodology, visualisation, writing – original draft, writing – review and editing.

CPS: Conceptualisation, writing-review/editing

ERG: methodology, supervision, funding acquisition, writing – review/editin

RMP: methodology, visualisation, funding acquisition, writing - review/editing.

PD: conceptualisation, formal analysis, project administration, resources, supervision, funding acquisition, writing – original draft and review/editing.

