## Supplementary information for "Investigating the evolutionary plasticity of influenza virus splicing using influenza A virus segment 7 as a model system"

### Supplementary Tables

| | Mutant | SD | SA | Amino acid changes in M1 | Amino acid changes in M2 |
|---|---|---|---|---|---|
| SD2 mutants | Weak (W) | 2 | Natural | - | - |
| | Medium (WT) | 2 | Natural | - | - |
| | Strong (S) | 2 | Natural | E8D and T9R | E8D and T9R |
| SD3 mutants | Weak (WT) | 3 | Natural | - | - |
| | Medium (M) | 3 | Natural | L42S | - |
| | Strong (S) | 3 | Natural | E40Q and L42S | - |
| | Strongest (S+) | 3 | Natural | E40Q and L42S | - |
| SD4 mutants | WT | No mRNA5 splicing detected | | - | - |
| | Strong SD4 | 4 | Natural | M165Q and T167S | - |
| | T167S | 4 | Natural | T167S | - |
| | T167S-no SD4 | No mRNA5 splicing detected | | T167S | - |
| SD5 mutants | WT | No mRNA6 splicing detected | | - | - |
| | Strong SD5 | 5 | Natural | L146Q and C148S | - |
| | L146Q | No mRNA6 splicing detected | | L146Q | - |
| | C148S | | | C148S | - |
| C870A-U872G (SA8) | WT | No mRNA7 splicing detected | | - | - |
| | C870A-U872G | 2 | 8 | - | - |
| SA7 mutants | WT | No SA7 usage detected | | - | - |
| | pY | | | E235F | - |

| | | | | | |
|---|---|---|---|---|---|
| | pY+A738G | 2 | 7 | E235F and Q238R | Δ aa 10-12 of M2 |
| | A738G-A749G | No SA7 usage detected | | Q238R and K242E | No M2 detected |
| | pY+A738G-A749G | 2 | 7 | E235F, Q238R and K242E | Δ aa 10-12 of M2 |
| SD4 + alternative SA mutants | Strong SD4-SA1 | 4 | 1 | M165Q, T167S, L237S and Q238E | Frameshift after T9 |
| | Strong SD4-SA2 | 4 | 2 | M165Q, T167S, N236T and L237V | Frameshift after T9 |
| | Strong SD4-SA3 | 4 | 3 | M165Q, T167S, E235A and N236D | Frameshift after T9 |
| | T167S-SA3 | 4 | 3 | T167S, E235A and N236D | Frameshift after T9 |
| SD3 + alternative SA mutants | SD3-SA1 | No splicing detected | | L237S and Q238E | No splicing detected |
| | SD3-SA2 | 3 | 2 | N236T and L237V | Ectodomain insertion See Fig 5E |
| | SD3-SA3 | 3 | 3 | E235A and N236D | Ectodomain insertion See Fig 5E |
| | SD3-SA4 | 3 | 4 | L237Q | Ectodomain insertion and substitution See Fig 5F |
| | SD3-SA5 | 3 | 5 | N236Q | Ectodomain insertion and substitution See Fig 5F |
| | SD3-SA6 | 3 | 6 | E235Q | Ectodomain insertion and substitution See Fig 5F |
| | SD3-SA7 | 3 | 7 | E235F, Q238R and K242E | Ectodomain deletion and substitution See Fig 5F |

**Supplementary Table 1.** Summary of non-synonymous changes in PR8 M1 and/or M2 resulting from changes to segment 7 splice sites, grouped by splice donor (SD) and splice acceptor (SA) site.

| Ion channel-related protein[1] | | Expression[2] | Titre[3] | Nucleotide substitution number | SD | SA |
|---|---|---|---|---|---|---|
| **M2 family** | | | | | | |
| M2 | 97 aa | +++++ | $10^8$ | - | 2 | Natural |
| Strong SD4-SA3 | 255 aa | ++++ | NR | 5 | 4 | 3 |
| SD3-SA2 | 130 aa | + | $10^2$ | 5 | 3 | 2 |
| SD3-SA3 | 131 aa | ++++ | $10^8$ | 5 | 3 | 3 |
| pY+A738G-A749G | 94 aa | ++ | $10^7$ | 5 | 2 | 7 |
| **M42 family** | | | | | | |
| M42 | 99 aa | +++ | $10^8$ | 2 | 3 | Natural |
| SD3-SA5 | 101 aa | ++ | $10^7$ | 4 | 3 | 5 |
| SD3-SA6 | 102 aa | + | $10^2$ | 4 | 3 | 6 |
| SD3-SA7 | 96 aa | ++ | $10^7$ | 7 | 3 | 7 |
| **Putative translation product from mRNA7** | | | | | | |
| M7 | 53 aa | Not detectable | $10^7$ | 2 | 2 | 8 |

Transmembrane domain
M1 ORF
M2 ORF (downstream of SA)
M42 ORF
Extra sequence because of splice site migration
Deletion because of splice site migration
Extra sequence because of removing stop codons
✕ Stop codon

**Supplementary Table 2. Phenotypic summary of PR8-based segment 7 with variant ion channel-related proteins produced by rewiring segment 7 splicing**. [1]Bars (approximately to scale) represent ion channel ORF structure, coloured according to the origin of the sequence. Overall ion channel length is also given. [2]Qualitative indication of relative expression level of the ion channel, defined by western blotting with an antiserum raised against M2 sequences downstream of the mRNA2 SD. [3]Approximate plaque titres.

A

| Mutant group | Mutant | Sequences | MaxEnt scores | Splice Site Strength |
|---|---|---|---|---|
| SD2 mutants | Weak | AAC\|CUACGU | 1.24 | |
| | Medium (WT) | AAC\|GUACGU | 9.51 | |
| | Strong | CAG\|GUACGU | 10.65 | |
| SD3 mutants | Weak (WT) | GAG\|GUUCUC | -1.72 | |
| | Medium | GAG\|GUUAGC | 3.77 | |
| | Strong | CAG\|GUUAGC | 4.9 | |
| | Strongest | CAG\|GUAAGU | 10.86 | |

B

| Mutant group | Mutant | Sequences | MaxEnt scores | Splice Site Strength |
|---|---|---|---|---|
| SD4 mutants | WT | AUG \| GUGACA | 0.78 | |
| | Strong SD4 | CAG \| GUAAGU | 10.86 | |
| | T167S | AUG \| GUAAGU | 11.01 | |
| | T167S no SD4 | AUG \| GUCUCC | -6.64 | |

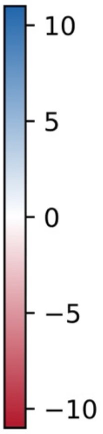


C

| Mutant group | Mutant | Sequences | MaxEnt scores | Splice Site Strength |
|---|---|---|---|---|
| SD5 mutants | WT | CUG \| GUAUGU | 8.14 | |
| | Strong SD5 | CAG \| GUAAGU | 9.80 | |
| | L146Q | CAG \| GUAUGU | 10.65 | |
| | C148S | CUG \| GUAAGU | 10.86 | |

D

| Mutant group | Mutant | Sequences | MaxEnt scores | Splice Site Strength |
|---|---|---|---|---|
| C870A-U872G | canonical SA | ATCTTCTTGAAA ATTTGCAG\|GCC | 6.29 | |
| | C870A-U872G | TTTTTCAAATGC ATTTACAG\|GCG | 7.68 | |

E

| Mutant group | Mutant | Sequences | MaxEnt scores | Splice Site Strength |
|---|---|---|---|---|
| Cryptic CAG in 746-748 | canonical SA (WT) | ATCTTCTTGAAA ATTTGCAG\|GCC | 6.29 | |
| | pY | ATCTTCTTTTTA ATTTGCAG\|GCC | 10.90 | |
| | A738G | AAAATTTGCGGG CCTATCAGAAA | 1.90 | |
| | pY+A738G | TTAATTTGCGGG CCTATCAG\|AAA | 3.28 | |
| | A738G-A749G | AAAATTTGCGGG CCTATCAGGAA | 3.14 | |
| | pYA738G-A749G | TTAATTTGCGGG CCTATCAG\|GAA | 4.53 | |

**Supplementary Figure 1. MaxEnt prediction scores for WT and mutant SD and SA motifs in PR8 Segment 7.** The mutational strategies and corresponding MaxEnt scores for (A) SD2, SD3, (B) SD4, (C), SD5 and canonical and cryptic SAs at (D) ntds 870-872 and (E) ntds 746-748 are shown. Mutations are underlined and the splice sites indicated by vertical lines. The heatmaps give a visual indication of the predicted splice site strengths.

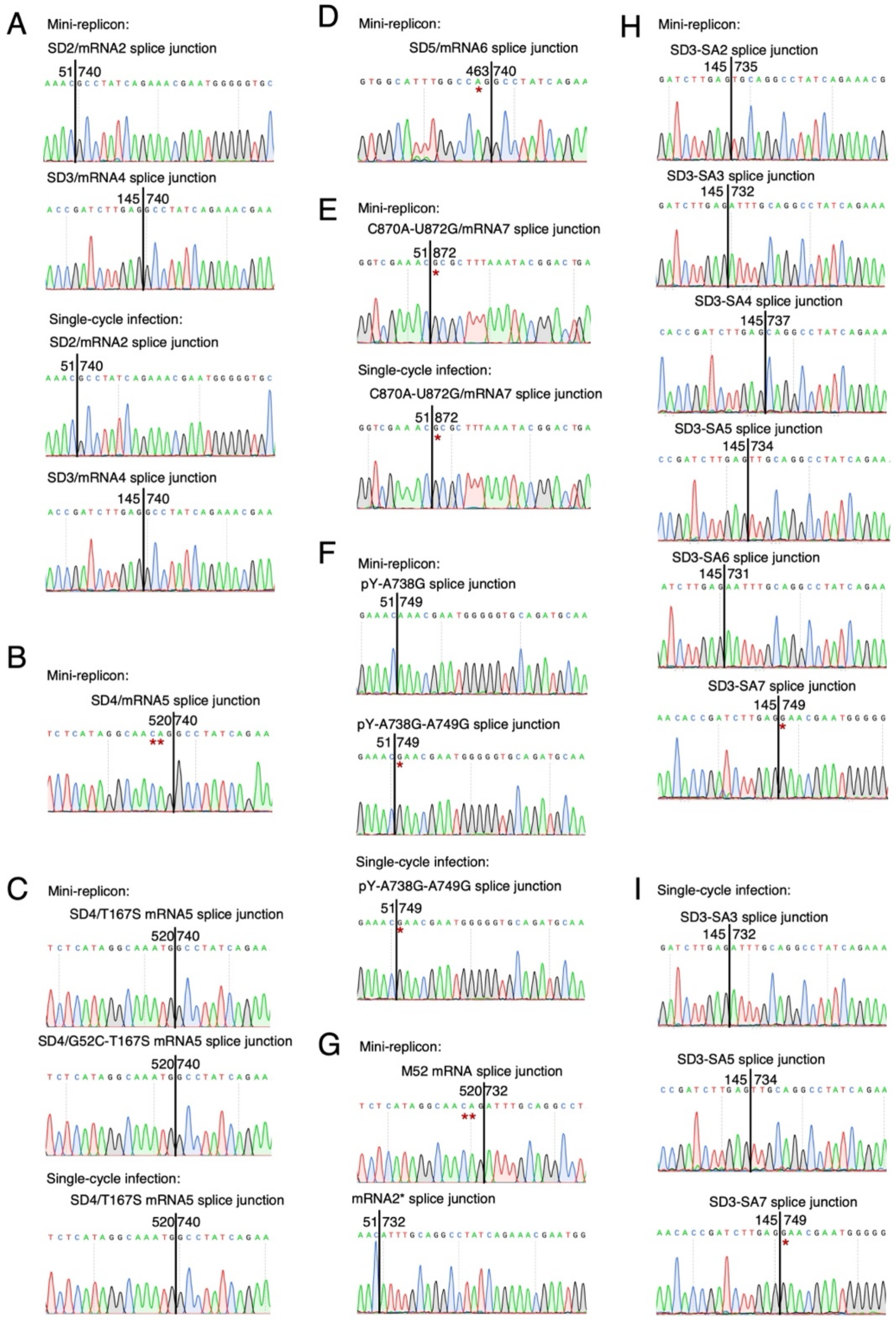


**Supplementary Figure 2. Sanger sequencing result for WT and novel spliced mRNAs from PR8 Segment 7.** Dye-deoxy sequencing traces from gel-extracted RT-PCR products are shown for the various spliced transcripts from mini-replicon assays or virus infections as labelled. Junction nucleotide coordinates are given. Asterisks indicate exonic mutations with respect to the WT PR8 sequence.

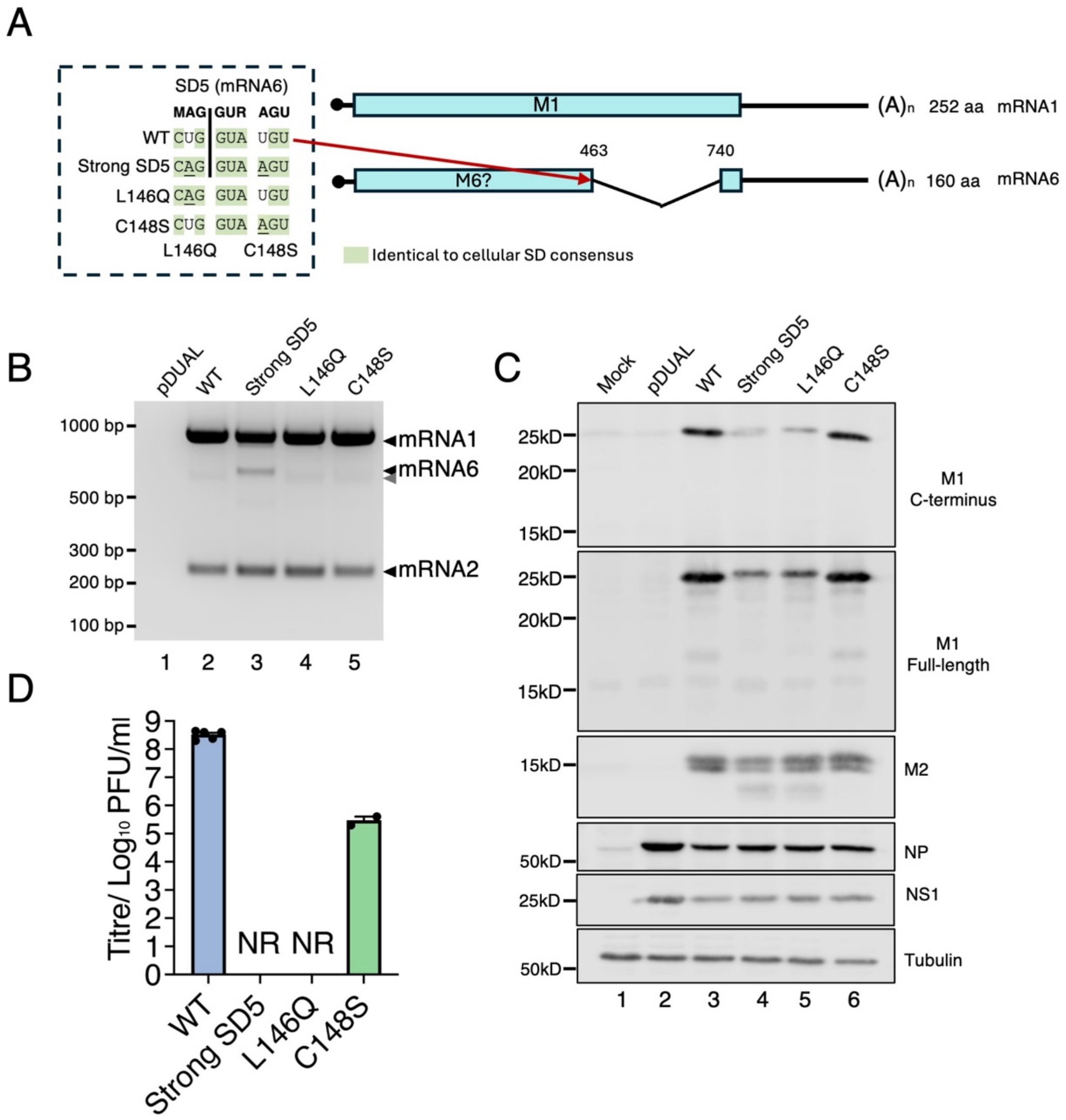


**Supplementary Figure 3. A virus with mRNA 6 is not viable.** (A) Mutational strategy for activating SD5/mRNA6 in PR8 segment 7. Identical nucleotides to the cellular SD consensus are shaded in green. (B, C) HEK 293T cells were co- or mock-transfected with IAV mini-replicon components, NS segment and the indicated segment 7 or pDUAL empty vector plasmids, harvested at 48h and analysed by (B) RT-PCR and agarose gel electrophoresis and (C) western blotting. The grey arrowhead indicates a probable heteroduplex of mRNA1 and 4 PCR products. (D) End-point titres at 48h p.i. of PR8 WT and mutant (except C148S) viruses grown in MDCK cells (MOI = 0.01). Data are presented as the mean ± SEM of at least two independent rescues. NR = not rescued. The titre of C148S was determined using the P0 stock.

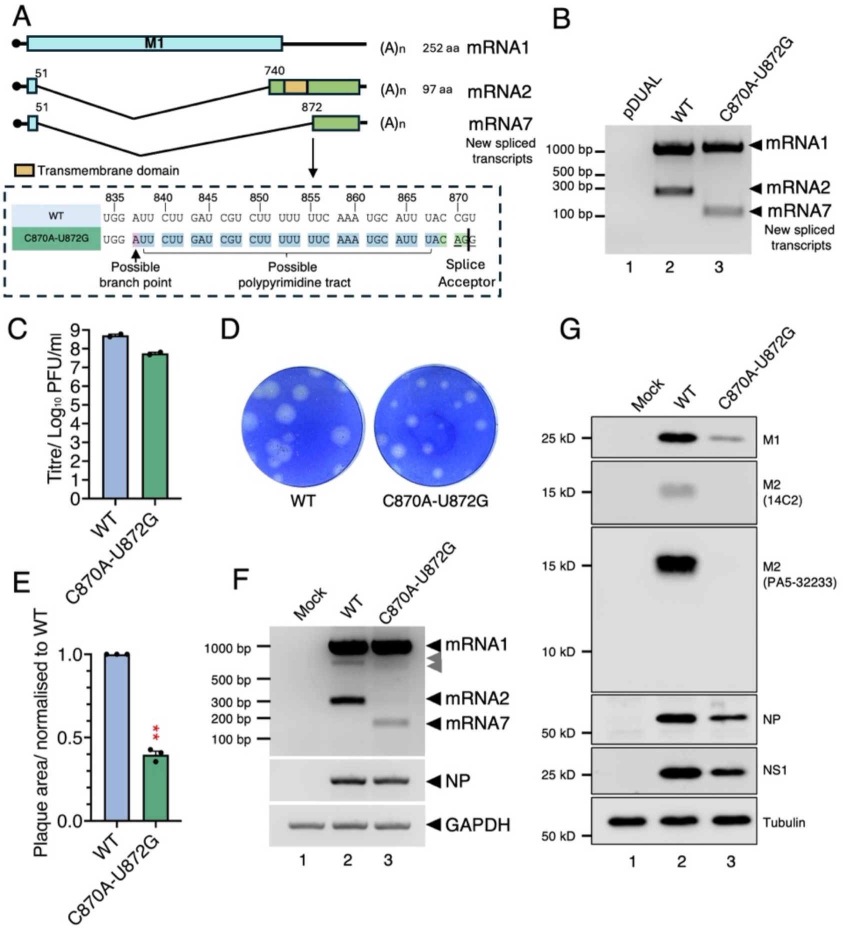


**Supplementary Figure 4. A new 3'-splice site can be activated by 2 ntd changes.** (A) Diagrammatic summary of a cryptic 3'-splice site in the M2 coding region. Upper section: the ORF structures of spliced and unspliced mRNAs are colour coded (blue, M1; light green, M2) and the transmembrane domain in M2 is highlighted in yellow. Lower section (dashed box): the nucleotide sequence of potential BP and PPT are highlighted in red and blue, respectively. The cryptic SA is highlighted in green. The mutation sites are underlined. (B) HEK 293T cells were co-transfected (or mock-transfected) with the mini-replicon components, NS segment and the indicated segment 7 or pDUAL empty vector plasmids, harvested at 48h and analysed by RT-PCR and agarose gel electrophoresis. (C) P1 stock titres of WT and mutant viruses. Data represent the mean of two independent rescues. (D) Representative images of plaque assays of WT and indicated mutants. (E) Quantification of the plaque size of PR8 WT and mutant viruses. Data are the mean ± SEM of 5-20 plaques normalised to the average WT value from at least three independent experiments. (F, G) MDCK cells were infected (or mock-infected) with the indicated viruses at an MOI of 5 and (F) total RNA isolated at 6 h p.i and the indicated mRNAs detected by RT-PCR or (G) lysed at 8 h p.i. and viral protein expression analysed by SDS-PAGE and western blotting. The grey arrowhead indicates a probable heteroduplex of mRNA1 and 2 or 7 PCR products.

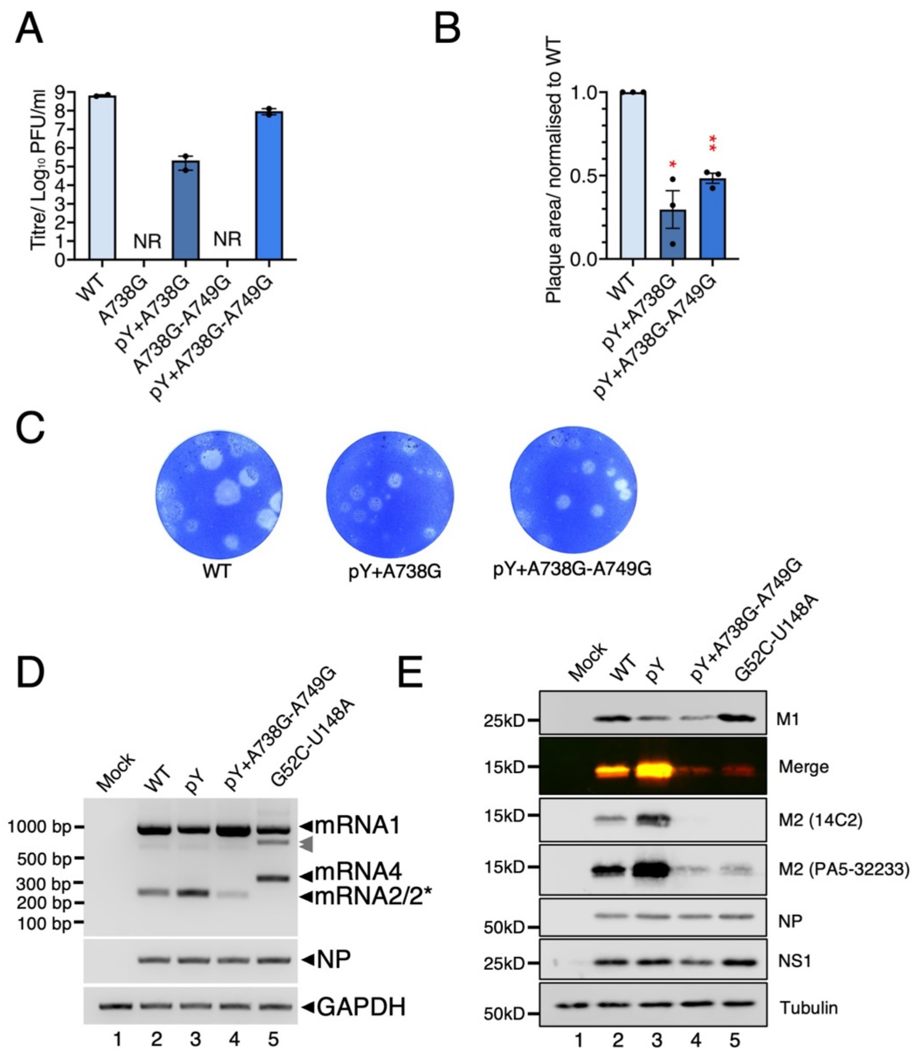


**Supplementary Figure 5. Viruses using the downstream SA are viable and produce a variant ion channel.** (A) P1 stock titres of WT and mutant viruses grown in MDCK cells. Data represent the mean of at least two independent rescues. (B) Quantification of the plaque size of PR8 WT and mutant viruses. Data are the mean ± SEM of 5-20 plaques normalised to the average WT value from three independent experiments. (C) Representative images of plaque assays of WT and indicated mutants. (D, E) MDCK cells were infected (or mock-infected) with the indicated viruses at an MOI of 5 and (D) total RNA isolated at 6 h p.i and the indicated mRNAs detected by RT-PCR, or (E) cells were lysed at 8 h p.i. and expression of the indicated proteins analysed by SDS-PAGE and western blotting. The grey arrowheads indicate a probable heteroduplex of mRNA1 and mRNA2/4 PCR products. * indicates co-migrating bands corresponding to the spliced products generated from cryptic SA usage.

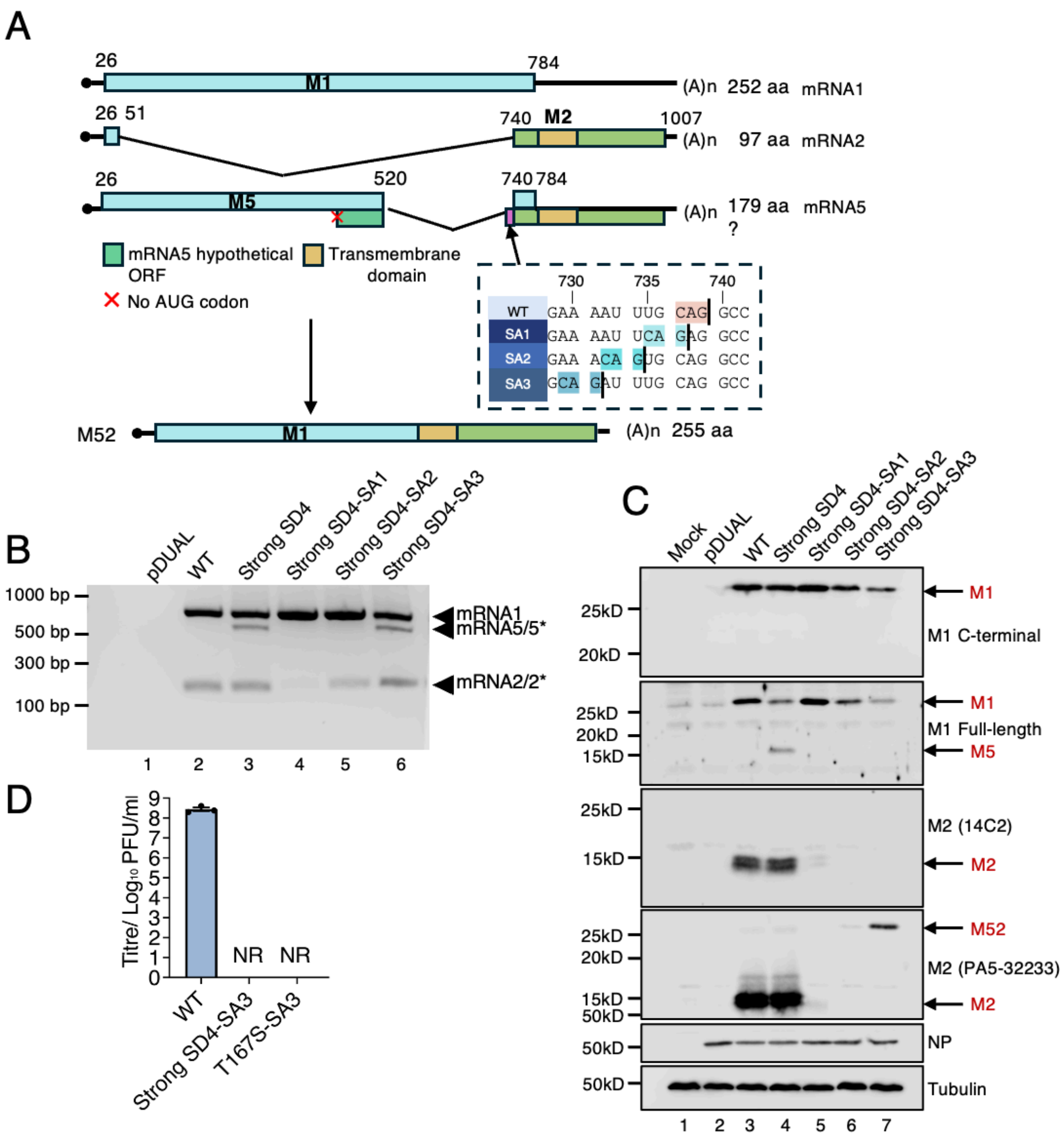


**Supplementary Figure 6. Rearranging segment 7 splicing to produce a novel M1-M2 fusion protein.** (A) Diagrammatic summary of mRNA splice variants. Potential ORFs are colour coded: blue, M1; light green, M2 and the M2 transmembrane domain is highlighted in yellow. Inset dashed box: canonical and novel SAs are highlighted in orange and blue shades respectively on WT and mutant sequences. (B, C) HEK 293T cells were co-transfected (or mock transfected) with the mini-replicon components, NS segment and the indicated segment 7 or pDUAL empty vector plasmids, then harvested at 48h and analysed by (B) RT-PCR and agarose gel electrophoresis and (C) western blotting as labelled. * indicates co-migrating bands corresponding to the spliced products generated from alternative SAs. M1, M2, M5 and 'M52' proteins are labelled by arrows respectively. (D) P1 stock titres of WT and mutant viruses grown in MDCK cells. Data represent the mean of at least two independent rescues.
